# Network Analyses of Brain Tumor Patients’ Multiomic Data Reveals Pharmacological Opportunities to Alter Cell State Transitions

**DOI:** 10.1101/2024.05.08.593202

**Authors:** Brandon Bumbaca, Marc R. Birtwistle, James M. Gallo

**Affiliations:** Department of Pharmaceutical Sciences, School of Pharmacy and Pharmaceutical Sciences, University at Buffalo, Buffalo NY, USA; Department of Chemical and Biomolecular Engineering, Clemson University, Clemson SC, USA; Department of Bioengineering, Clemson University, Clemson SC, USA

## Abstract

Glioblastoma Multiforme (GBM) remains a particularly difficult cancer to treat, and survival outcomes remain poor. In addition to the lack of dedicated drug discovery programs for GBM, extensive intratumor heterogeneity and epigenetic plasticity related to cell-state transitions are major roadblocks to successful drug therapy in GBM. To study these phenomenon, publicly available snRNAseq and bulk RNAseq data from patient samples were used to categorize cells from patients into four cell states (i.e. phenotypes), namely: (i) neural progenitor-like (NPC-like), (ii) oligodendrocyte progenitor-like (OPC-like), (iii) astrocyte-like (AC-like), and (iv) mesenchymal-like (MES-like). Patients were subsequently grouped into subpopulations based on which cell-state was the most dominant in their respective tumor. By incorporating phosphoproteomic measurements from the same patients, a protein-protein interaction network (PPIN) was constructed for each cell state. These four-cell state PPINs were pooled to form a single Boolean network that was used for *in silico* protein knockout simulations to investigate mechanisms that either promote or prevent cell state transitions. Simulation results were input into a boosted tree machine learning model which predicted the cell states or phenotypes of GBM patients from an independent public data source, the Glioma Longitudinal Analysis (GLASS) Consortium. Combining the simulation results and the machine learning predictions, we generated hypotheses for clinically relevant causal mechanisms of cell state transitions. For example, the transcription factor TFAP2A can be seen to promote a transition from the NPC-like to the MES-like state. Such protein nodes and the associated signaling pathways provide potential drug targets that can be further tested *in vitro* and support cell state-directed (CSD) therapy.

## INTRODUCTION

Glioblastoma multiforme (GBM) is a highly aggressive brain cancer and little progress has been made in improving survival outcomes. Temozolomide (TMZ), a DNA alkylating agent, is the primary standard of care drug along with radiotherapy (RT) but has only resulted in a median survival of 15 months and a 5-year survival rate of 5% [1, 2]. Further attempts at improving patient outcomes have been fruitless. Bevacizumab, a monoclonal antibody which targets VEGFA, was approved for the treatment of recurrent GBM (rGBM) but has failed to achieve significant improvements in overall survival (OS) [3]. Both erlotinib and gefitinib have failed to improve patient outcomes in GBM, despite being EGFR inhibitors designed for patients with mutated EGFR [4, 5]. Immune checkpoint inhibitors have fared poorly as well, as both nivolumab and pembrolizumab were ineffective in patients with rGBM [6, 7]. The reasons for treatment failure are multi-faceted and intratumoral heterogeneity (ITH) and cellular plasticity are two phenomena that are known to contribute to drug resistance [8–11].

ITH in GBM has been explored extensively and multiple classifications have emerged for interpatient classification of GBM subtypes. Bulk transcriptomic analyses, for example, have been used to identify three main molecular subtypes of GBM across patients: proneural, mesenchymal, and classical [12–14]. Other works have focused on common genetic alterations, such as the EGFRvIII mutation, EGFR copy number amplification, TP53 deletion, and NF1 alterations [11, 15–17]. Mutant isocitrate dehydrogenase 1 (mIDH1) has become a distinct marker of prolonged survival in GBM as patients with this mutation have a median survival of 31 months [18, 19]. Additionally, the methylation status of O6-methylguanine-DNA-methyltransferase (MGMT) is a valuable prognostic for TMZ efficacy [20, 21]. All these events may be associated with molecular subtypes and clinical outcomes [11, 15] and more work is needed to understand these relationships in detail.

More recent research has given insight into the epigenetic plasticity of GBM tumor cells and the inherent heterogeneity within a single tumor. A high amount of stemness increases plasticity of the tumor and is positively associated with tumor grade [22]. This plasticity is thought to drive tumor self-renewal and therapeutic resistance through the existence of multiple phenotypes or cell states and transitions [23–27]. A recent effort by Neftel and colleagues [28] utilized single cell sequencing (scRNA-seq) to identify four primary cell states for GBM (Mesenchymal-like, Astrocyte-like, Neural-progenitor-like, and Oligodendrocyte-like) and showed how it is possible for these cell states to transition to one another. Another work [29] also tracked changes in single cell populations of GBM cells *in vitro* in response to TMZ exposure, showing how drugs may induce transitions and alter the phenotypic landscape of GBM.

Since drugs can induce cell state transitions, it may be possible to predict and control these transitions if their mechanisms are understood. For example, one treatment strategy proposed by Prager et al. [23] is to use drugs to shift the GBM cell population into one attractor or phenotypic state and dose with another drug which is highly effective against that particular phenotype. Indeed, controlling cell state transitions has been of interest since C.H. Waddington first proposed his landscape of epigenetics [30]. Rukhlenko et al. [31] showed it is possible to develop models which can describe phenotypic changes and proposed mechanisms on how to control these changes. However, these mechanisms are not yet fully understood in GBM.

Our lab has previously explored how epigenetics and cell state transitions may result in TMZ resistance in GBM, and how it may be possible to overcome this resistance via cell state-directed (CSD) therapy [32, 33]. In this work, we utilized a multi-step computational pipeline that incorporated multi-omics data from GBM patients to more broadly explore mechanisms for cell-state transitions and identify key genetic and epigenetic mediators. First, for each patient the dominant cell state defined by Neftel and colleagues was determined using single nucleus RNA sequencing (snRNA-seq) in a subpopulation or reference dataset and bulk RNASeq data from the remaining patients by deconvolution. Next, by incorporating phosphoproteomic data for all patients, PPINs were built for each of the four cell state groups. These four PPINs were subsequently combined to create a single Boolean network capable of describing the relationships between the four cell states. We then used *in silico* simulations to propose testable pharmacological interventions to either promote or prevent cell state transitions. Finally, the results from these simulations were used as input to a machine learning model capable of predicting the dominant cell state of another independent clinical GBM dataset. By combining simulation results and the predictive machine learning model, we propose plausible causal mechanisms of action of cell state transitions in GBM patients that may form the basis for pharmacological targeting and CSD therapy.

## RESULTS

### Generating Protein-Protein Interaction Networks for each Cell State

Although every GBM tumor has multiple cell states within it, most tumors have a dominant cell state that is highest in proportion. We hypothesized that given data for enough patients that are dominant for the same cell state, even if bulk data, we could extract sufficient information to propose protein-protein interaction networks that are enriched for cell-state specific features corresponding to this dominant cell state. As a first step towards exploring this hypothesis, a multi-step pipeline – see Figure 1 - was used based on a suite of publicly available data (Wang data set, described next) that yielded unique PPINs for each GBM cell state (based on the Neftel definitions). The Wang data set includes 100 GBM patients consisting of a collection of multiomics measurements and clinical information from each patient. Our work utilizes 92 of these samples that are from wtIDH1 GBM patients (the remaining 8 were mutant IDH1 samples and excluded), and incorporates bulk RNAseq, phosphoproteomics, and single nucleus RNA sequencing (snRNAseq). The results of this pipeline are described below and the analytical details in the Methods section.

**Figure 1.**
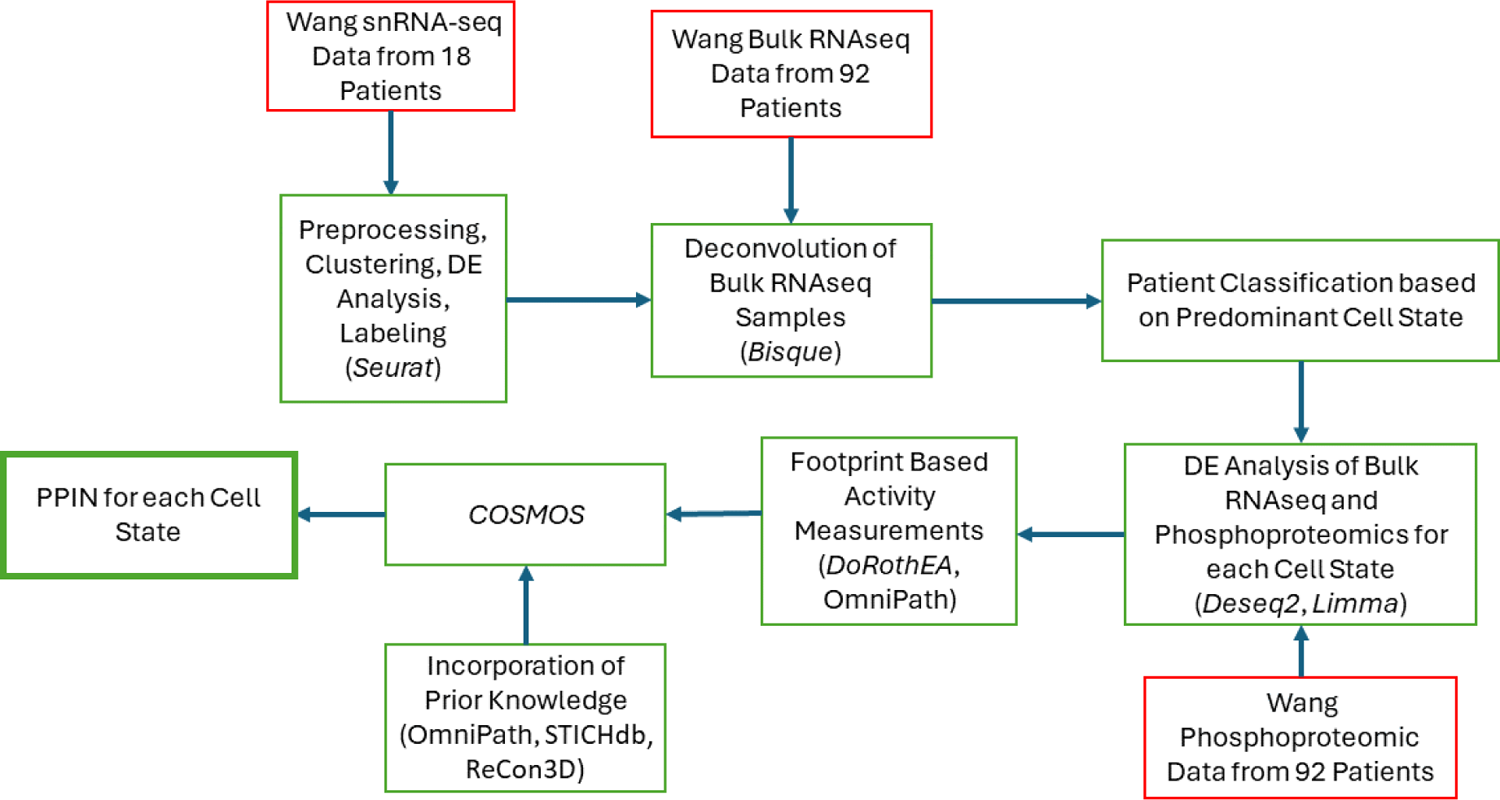
Workflow for the creation of PPINs. All omics data (red box) were accessed via the CPTAC portal (https://proteomics.cancer.gov/data-portal) and originally published by Wang et al. 2021 [34]. The software used in each step (packages/databases) are noted in each green box in parentheses. *Seurat v4* [35], *Bisque* [36], *limma* [37], and *Deseq2* [38], were used for snRNA-seq analysis, deconvolution, phosphoproteomic analysis, and bulk RNAseq analysis, respectively. Activity inferences for transcription factors and kinases were estimated using *DoRothEA, OmniPath,* and *DecoupleR,* while prior knowledge was combined from three databases (*STICHdb*, *ReCon3D*, and *OmniPath*).

### Single Cell Analysis

The Wang snRNAseq data was designated as the reference data to label cells based on the Neftel cell types. The 18 snRNAseq samples from wtIDH1 GBM patients were imported into the R package *Seurat* and analyzed as described in the methods section. In brief, cells were analyzed for quality and clustered based on similarity of gene expression, resulting in forty clusters. These forty clusters were tested for enrichment (hypergeometric test) of gene modules for Mesenchymal-like (MES), Astrocyte-like (AC), Neural-progenitor-like (NPC), and Oligodendrocyte-like (OPC) cell states along with other healthy cell types such as astrocytes (Astro), neurons, and oligodendrocytes (oligo) The enrichment of all cell clusters can be found in the supplementary material (S1). Each cluster was labeled with the most enriched gene module and all the clusters were visualized together via UMAP (S2).

### Deconvolution and Patient Classification

Once all the clusters were labeled with the most enriched cell state, each individual cell in each cluster is labeled with the same cell state as that cluster. It is therefore simple to determine the predominant cell state for those 18 patients for whom there are snRNA-seq samples available, as each cell is marked with the original patient IDs. However, there are an additional 74 patients in the Wang dataset which do not have snRNAseq samples but do have bulk RNAseq samples. To get an estimate of the cell composition of all patient samples, the R package *Bisque* was used to deconvolve bulk samples into individual cell type estimates (Figure 2a) [36]. These analyses suggest that each patient predominantly expresses one of the four major cell types (MES, AC, NPC, OPC). We then grouped the patients together based on their most dominant cell state. Figure 2b shows the results of the patient classification with most patients predominantly expressing either the MES-like or NPC-like cell state.

**Figure 2.**
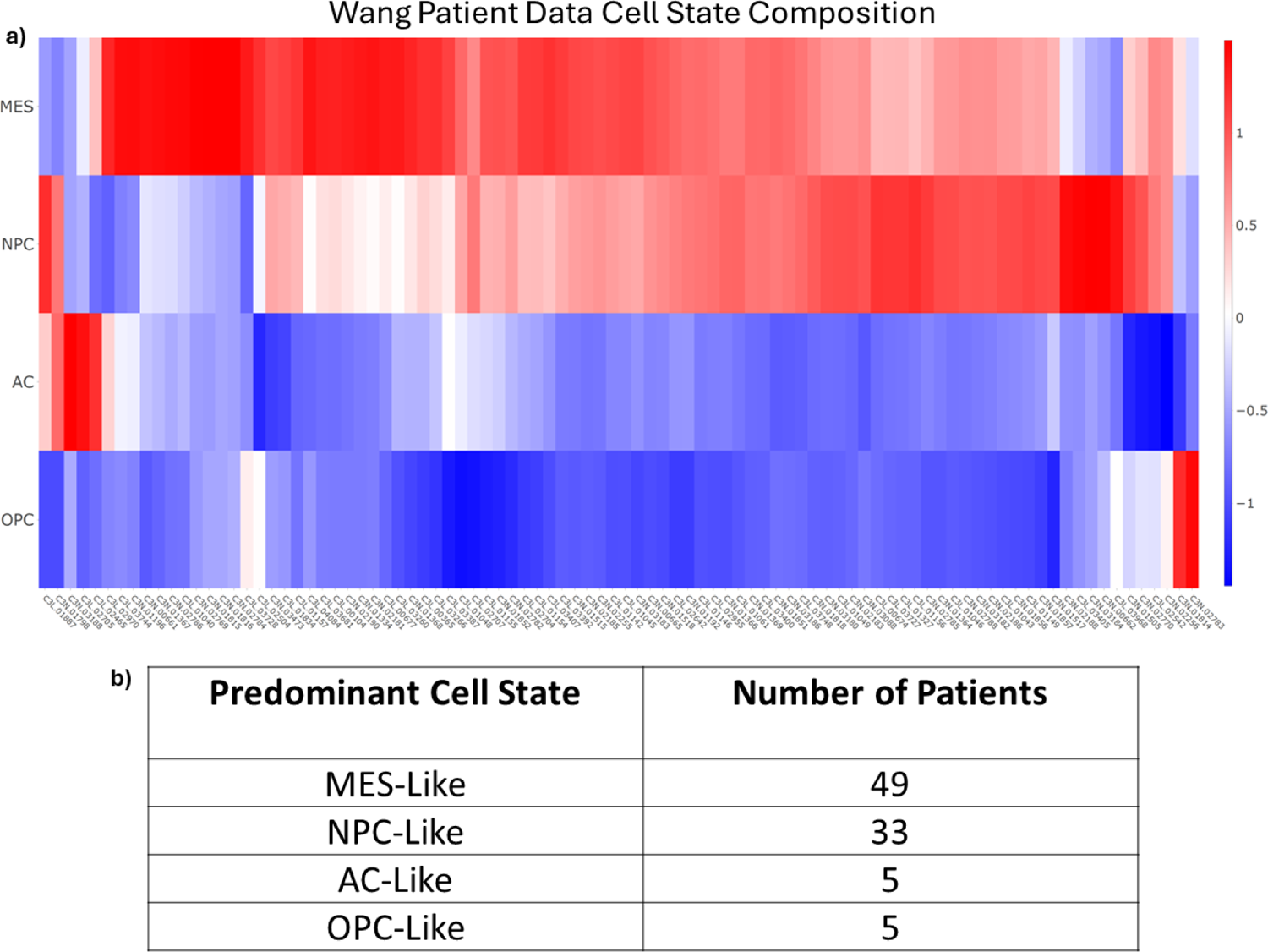
Bisque Deconvolution. **a)** The deconvolution results of the bulk RNAseq data from Wang [34]. Rows are the four cell states and columns are the patient IDs (as used in Wang). Patients were subsequently binned into four groups based on the predominant cell state. **b)** The total number of patients binned into each of the four cell states.

### Cell-State Specific and Integrated Networks

Patients were divided into the four cell state groups based on their predominant cell state following the deconvolution analysis. Four PPINs were constructed using these patient subgroups and the COSMOS pipeline (see methods section) [39]. In brief, bulk RNAseq and phosphoproteomics for each subpopulation were analyzed for differentially expressed entities. The activity of both transcription factors and kinases were estimated using the WMEAN algorithm from the *decoupleR* R package [40]. These activity measurements are then used to identify the most likely interactions among all proteins and create the PPINs [39]. On average, the networks have 65 nodes and 122 edges (S4, a-d).

We were curious as to how similar or different the PPINs were to each other. Using the 50 hallmark pathways from the Human Molecular Signatures Database (MSigDB) as a reference, we found that all the cell states included a high number of G2M checkpoint pathway proteins (Table 1) [41]. Perhaps surprisingly, all cell state PPINs contain many genes from the allograft rejection pathway. However, this pathway contains genes from common growth and proliferative pathways including *EGFR* and *AKT1*, which may explain its presence in the networks. Additionally, the AC, MES, and NPC cell states contained many proteins from the PI3K/AKT signaling pathway, but the OPC state did not. The AC state contained the least number of proteins from the TNFA/NFKB signaling pathway. The OPC and NPC states are upregulated for the interferon gamma and JAK/STAT pathways, but the MES and AC states are not. These four networks were subsequently merged to form the final PPIN (S4, e). All unique interactions across the four networks are included for a total of 144 proteins and 380 interactions.

**Table 1.** Hallmark Pathways in each Cell State. The number of proteins present from the top 10 most represented hallmark pathways in the MSigDB in each of the four-cell state PPINs.

| Pathway/Cell State | AC | MES | NPC | OPC |
| --- | --- | --- | --- | --- |
| G2M CHECKPOINT | 8 | 7 | 11 | 7 |
| PI3K AKT MTOR SIGNALING | 9 | 10 | 7 | 5 |
| TNFA SIGNALING VIA NFKB | 3 | 7 | 11 | 7 |
| ALLOGRAFT REJECTION | 7 | 7 | 6 | 6 |
| IL6 JAK STAT3 SIGNALING | 2 | 5 | 6 | 7 |
| INTERFERON GAMMA RESPONSE | 2 | 4 | 7 | 6 |
| UV RESPONSE DN | 5 | 5 | 5 | 4 |
| HYPOXIA | 3 | 4 | 5 | 6 |
| APOPTOSIS | 4 | 5 | 5 | 4 |
| E2F TARGETS | 4 | 4 | 6 | 4 |

### Steady-State Analysis

The final PPIN was converted into a Boolean network to enable simulations to steady-state and to perform in silico protein knockout simulations (next section). In this construction, each node may be in one of two states at a given time: “ON” (1) or “OFF” (0). A critical component of the conversion from a static PPIN to a Boolean model is generating the logical equations of the network. To accomplish this task, we aimed to match the steady-state of each node in the network to its state as defined by the COSMOS PPIN. In other words, when run to steady-state each node in the model should be either active or inactive. An active node is equivalent to an upregulated node in COSMOS, while an inactive node is equivalent to a downregulated node. By conducting this analysis for all four cell states, we can effectively define the logical rules that govern the network. MaBoSS (Version 2.0) [42] was used to perform the steady-state simulations and create logical rules capable of achieving this goal. The full list of rules can be found in the supplemental material (S3).

We then sought to compare the four different steady-states (one for each cell state) of the Boolean network. Using MaBoSS, the network was run to steady-state starting from each of the predominant cell states (four times) and the results were compared (Figure 3b). A similarity score was defined as the percentage of nodes in the same “ON” or “OFF” state as at time zero. Notably, the MES and AC state were the most similar indicating greater overlapping activity. Interestingly, there are some differences in steady-state similarity depending on which cell state is the initial starting point, indicating non-linear responses within the network.

**Figure 3.**
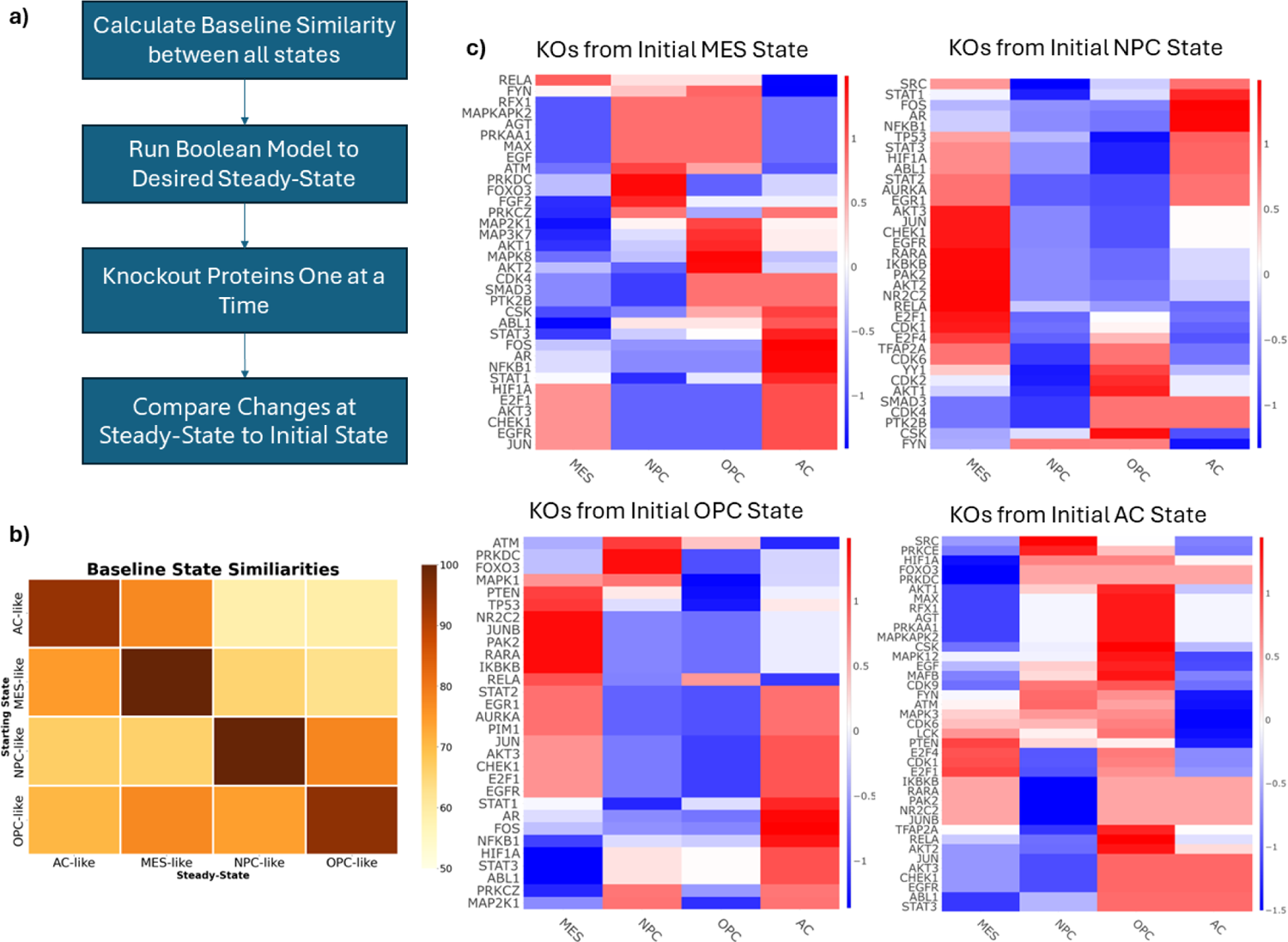
Boolean Simulations. **a).** Workflow for the Boolean Simulations. **b)** The similarity between each cell state at steady-state. Similarity is defined as the percentage of nodes in the same “ON” or “OFF” state at steady-state compared to its initial state in the Boolean network. **c)** Z-normalized Hamming Distances of the knockout simulations conducted with the Boolean Network. Positive scores (red) indicate a knockout shifted the network toward that cell state, while negative scores (blue) indicate a shift away from a cell state. Each row represents an individual protein KO, and each column describes the changes with respect to each cell state. Each heatmap represents the results from starting from each of the cell states. Starting at the top left and going clockwise: MES, NPC, OPC, AC.

### *In Silico* Knockout Simulations

Starting from the steady-state of each predominant cell state, nodes were knocked out one at a time and the resulting state was compared to the original steady-state of the network. This was conducted for each active node in each cell state. The Hamming distance was calculated for each simulation and the z-normalized distances are shown in Figure 3c. Notably, knocking out *TP53* or *PTEN* (two commonly mutated proteins in GBM [43, 44]) drives the network toward the MES state and may be a key cause of resistance to TMZ and RT [14, 45, 46] Interestingly, *STAT3* knockout drives the network away from the MES state, toward the AC state. The transcription factors E2F1 and E2F4 are predicted to be critical for maintaining the NPC-like state.

### Machine Learning Models Predict Cell State of GBM Patients

Given that our Boolean model was constructed from patient data (i.e. Wang), we sought to determine if our model simulations are representative of other patient samples as well. Our simulations suggest that many different perturbations could cause a change in cell state, but it remains unclear which of these perturbations, if any, may occur in patients. We developed a workflow (Figure 4) to help clarify these two questions. Using the KO model simulations from the Wang data as input (Figure 3c), we trained four different machine learning models, namely: multinomial logistic regression, K-nearest neighbor (KNN), random forest (RF), and boosted trees (XGBoost). The Glioma Longitudinal Analysis (GLASS) Consortium has created a database of primary and recurrent glioma tumors including clinical characteristics, bulk RNAseq and other omics data, and is an ideal independent dataset to validate that our model predictions are accurately capturing clinical cell states. We used bulk RNAseq data from 201 patients. Just as with the Wang dataset, only wtIDH1 samples were considered. The data was processed to match the format of the Boolean model. Briefly, bulk RNAseq were deconvoluted to determine the predominant cell state of each patient. Differentially expressed genes were identified for each sample and used to estimate transcription factor activity. In lieu of phosphoproteomic data, we derived a relationship between RNAseq and phosphoproteomic activity measurements using the Wang dataset as reference (see Methods). The activity measurements are then converted to Boolean values. These values are the input to the ML models as our test cases. We tested the models’ abilities to correctly predict the predominant cell state of the GLASS dataset. Primary and recurrent tumors were tested independently, and additional datasets adding noise into the activity measurements were also tested.

**Figure 4.**
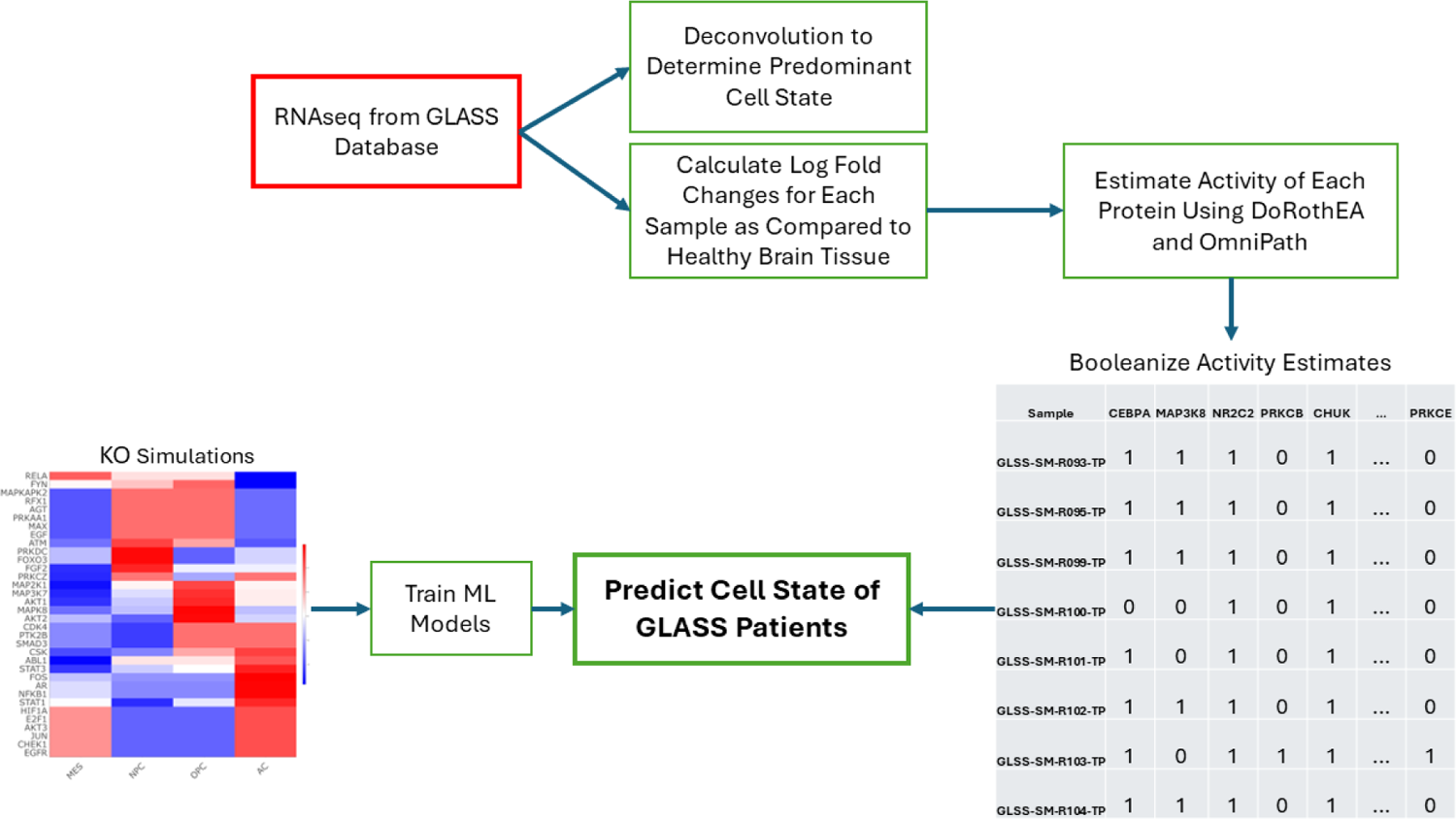
Machine Learning Workflow. Protein KO simulations are used to train machine learning models to predict cell state. RNAseq data from the GLASS database were used to determine the predominant cell state for each GLASS sample. These data were compared to healthy brain tissue to determine differentially expressed genes and estimate protein activity. These protein activities are booleanized to match the format of the Boolean simulations and become the test cases for machine learning performance.

XGBoost performed the best out of all the machine learning models for every dataset. When breaking down the full results by cell state and tumor type (primary or recurrent), we considered the balanced accuracy of each cell state as our objective metric, due to the massive class imbalances present in the dataset. The MES and NPC primary tumor cell states were predicted with a balanced accuracy of 99.1% and 92.3%, respectively. The recurrent tumors of the same cell states were predicted with a balanced accuracy of 98.2% and 89.8%, respectively. The primary tumors of the AC and OPC cell states were predicted with a balanced accuracy of 45.6% and 50%, while the recurrent tumors were predicted with a balanced accuracy of 41.6% and 50% respectively. All the results from the four different ML models can be found in the supplemental material (S5). Given the MES and NPC cell states are the two most common predominant states, our model performed well overall, however the results indicate that our simulations for the AC and OPC state may not be capturing clinically relevant phenomena. More work is needed to understand how the model may be misinformed.

### XGBoost Interpretations Provide Mechanistic Hypotheses for Cell-State Transitions

Historically, ensemble methods of decision trees have been difficult to interpret. However recent advances in machine learning techniques have opened the door for interpreting these traditionally “black box” models. One such method is Shapley Additive Explanations (SHAP) which uses game theory to understand the contribution of each feature to the final model predictions (Figure 5b) [47]. While feature contributions for all states are calculated, we focused on the MES and NPC states as our model predicted these cell states well. Importantly, *CDK2, HGF, JUNB, IRF1, FGF2, and TGFB1* are identified as important predictors of the MES and NPC states.

**Figure 5.**
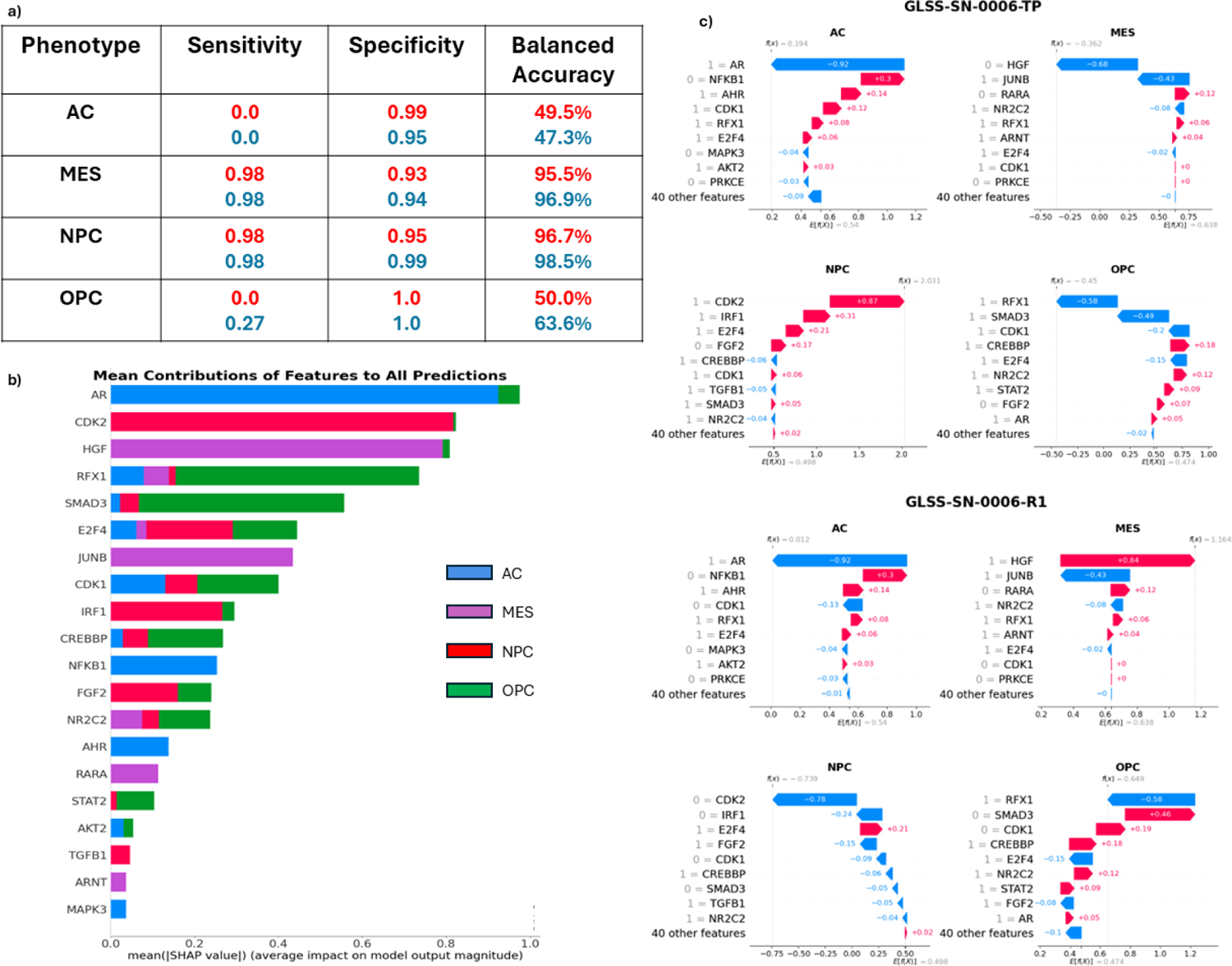
Machine Learning Results. **a)** XGBoost Performance. The table shows the results of XGBoost predictions separated by phenotype and tumor type. **b) The Mean contribution of each Feature to all Cell State Predictions**. Shapley values were calculated for each of the 201 tumors and the average score is shown here. Each row represents the individual gene (feature) contribution to each of the 4 cell states and shown as a sum **c) Shapley Plots of a Primary and Recurrent Tumor of One Patient.** Each primary and recurrent (22 months later) tumor has four Shapley plots, one for each of the four cell states. Each plot depicts how the most critical proteins affects the prediciton of each cell state from the machine laearning model. For this patient, the primary tumor was correctly identified as an NPC-like tumor (2^nd^ row, left) with CDK2 the most contributing positive feature, while the recurrent tumor was correctly identified as an MES-like tumor (3^rd^ row, right) with HGF the most contributing positive feature.

One additional benefit of the Shapley approach is that each individual patient sample can be investigated for which features contributed to the predicted cell state. Specifically, the method could indicate what features (genes) probabilistically are involved in a predicted cell-state transition (if a transition occurs). To illustrate this approach, we will use one example from a specific patient in the GLASS dataset who had both a primary and a recurrent tumor sample. As seen in Figure 5c, this patient’s primary tumor was correctly classified as a predominately NPC-like tumor. In this case, *CDK2* and *IRF1* being in the active state were the largest contributors to correctly identifying the cell state of the tumor as NPC. On the other hand, *HGF* being in the inactive state was a major contributor to why XGBoost did not choose the MES state as the predominant state. Subsequently, this patient underwent standard of care therapy (TMZ and RT), and another tumor sample was taken 22 months later (Figure 5c). In this tumor, we again correctly identified the predominant cell state, MES. Using the SHAP values we can see what key proteins changed state over the course of therapy, and at the very least were associated with the shift from NPC to MES. In this case, both *CDK2* and *HGF* changed states. *CDK2* became less active while *HGF* became more active indicating a switch to a MES-like tumor.

### Combining SHAP Interpretations with Knock Out Simulation Results

The protein knockout simulations provided a plethora of potential perturbations which could cause a cell-state transition, but did not provide information on which of these transitions may be most likely to occur in a clinical setting. On the other hand, our machine learning model was able to correctly identify the key proteins that switched state over the course of therapy, but do not consider the perturbations or dynamics that may have led to these changes. We wanted to combine these two pieces of information to create causal hypotheses of cell-state transitions. We have already shown that high *CDK2* and *HGF* activity are positively correlated to the NPC and MES state, respectively, and that the inactivation of *CDK2* and activation of *HGF* in response to standard of care are clincially relevant changes indicative of a cell-state transition from the NPC to the MES state (figure 5c). However, the question still remains as to how CDK2 and HGF become inactive and active. In other words, what are the transcriptional or signaling changes that are capable of bringing about significant decreases in *CDK2* activity or increases in *HGF* activity? We can use our protein KO simulations from the NPC state to search for answers to this question. Looking at figure 3c, we can observe that many KOs from the NPC may shift the cell state toward the MES state. However, by focusing on the HGF pathway we found that a KO of transcription factor *TFAP2A* is capable of causing *HGF* activation. *TFAP2A* activates *PTEN*, which is a known inhibitor of *HGF* and *HGF* signaling Figure 6). We thus can connect *TFAP2A* silencing with *HGF* actviation and and a transition from the NPC-like to the MES-like state. Another study has corroborated this mechanism of action, showing that *TFAP2A* transcriptional binding sites are hypermethylated in recurrent GBM tumors following treatment from standard of care [48]. Threrefore, the epigenetic silencing of *TFAP2A* is a plausible hypothesis for cell states transitions from the NPC to the MES state in GBM. Decitabine, a hypomethylating agent, has been shown to increase *TFAP2A* expression/activity in GBM and increase PD-L1 expression [49]. The effect of decitabine on PTEN expression GBM has not been observed, however it has increased PTEN expression in other cell lines [50, 51]. Increasing PTEN activity with decitabine treatment may help prevent the MES state and be an effective method of CSD therapy.

**Figure 6.**
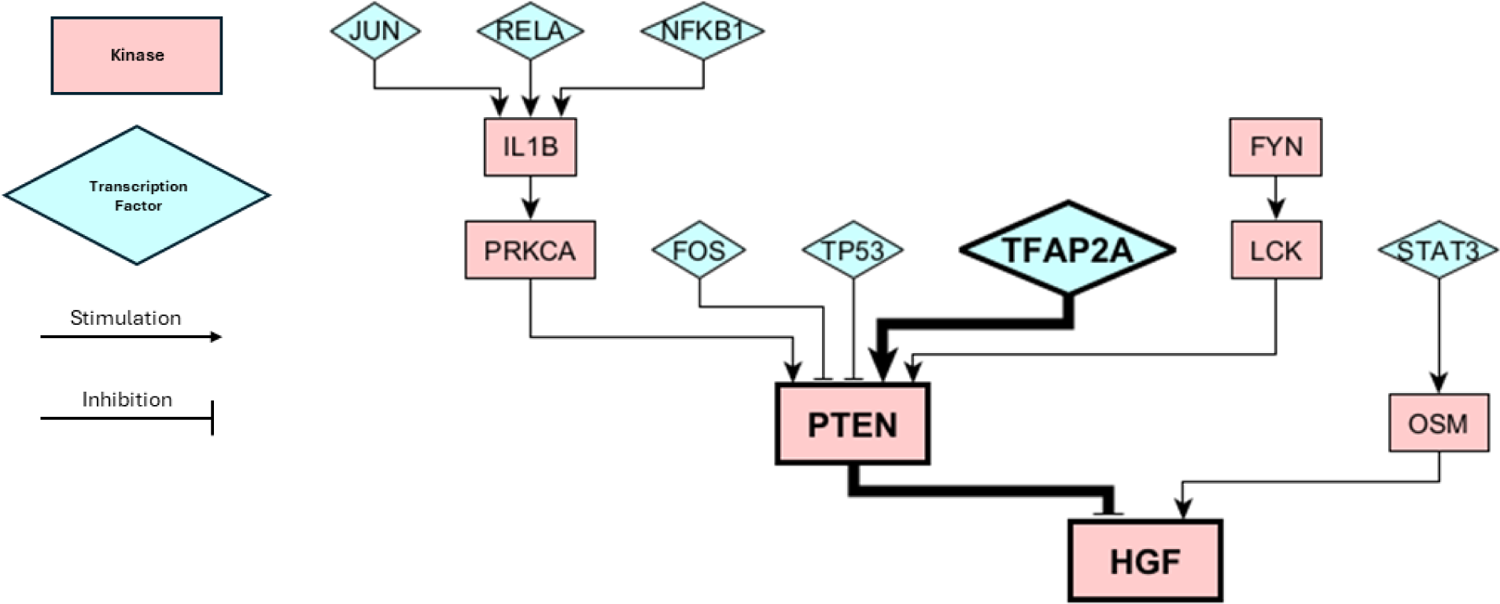
Submodule of our Boolean Network. The bolded proteins show the pathway involved in causing a cell-state transition from NPC to MES, as predicted by our knockout simulations and XGBoost predictions. Kinases are depicted by pink rectangles and transcription faactors are depicted as blue triangles. Activating/stimulating interactions are depicted by arrows and inactivating/inhibitory interactions are depicted with T-bars.

## DISCUSSION

There remains a critical need to improve the treatment of GBM. Due to the high degree of heterogeneity and plasticity of GBM cells, controlling cell state transitions may be a viable treatment strategy, and previously referred to as cell state-directed (CSD) therapy [33]. There have been a few different classifications proposed to describe the cell states of GBM [12, 13, 52]. These classifications utilize different types of omics data. We chose the Neftel et al. classifications as these phenotypes are defined at the single cell level and are robust across multiple different datasets [53–55]. While other investigations have explored the transcriptional differences between GBM cell states, this is the first work to incorporate phosphoproteomic measurements into PPINs for a relatively large number of GBM patients (92). The combined PPIN represented four GBM cell states considered relevant to maintenance of tumor growth and treatment resistance. The combination of phosphoproteomics with RNAseq data provides not only a more robust network that connects gene regulatory networks and downstream signaling pathways but also possible protein targets amenable to pharmacological intervention. This network model could be a foundation to implement CSD therapy to either promote or prevent desired cell state transitions [33]. To that end, a Boolean network model derived from the combined PPIN was constructed and used to conduct *in-silico* protein knockout simulations that serve as proxies for pharmacological intervention. Finally, our machine learning model was able to identify the relationships between the activity of specific proteins and cell states to predict phenotypes in clinical samples. Connecting these predictions with our simulations provided a more complete robust picture of cell state transitions and more nuanced hypotheses.

Although the Boolean modeling approach is advantageous for large-scale pharmacodynamic simulations, it can be appreciated that converting the current model into a biochemical mechanistic model may be rewarding. Nonetheless, the current analyses have generated viable hypotheses from large multi-omics GBM patient datasets. The model simulations and ensuing machine learning analyses led to testable hypotheses of how pharmacological intervention of protein targets could prevent or promote a cell state transition. It will be important to associate these or other quantifiable cell states with aggressive cell proliferation and treatment resistance to fully implement CSD therapy.

There are limitations to this work. First, the PPIN model only includes intra-cellular signaling networks whilst it is known that other cell types exist in the tumor microenvironment including immune cells that affect the phenotypic behavior of cancer cells [56–59]. Such cell-cell interactions are ripe for future exploration and the multi-omics pipeline could be adapted to develop GBM cell and microenvironment interactions and ultimately a revised PPIN [60]. A second limitation is that the model was built with data mostly reliant on bulk tumor samples representing a conglomerate of cell populations. Given what is known about ITH in GBM and shown in this work, it is certain that no one tumor will ever express only one phenotype and single cell multi-omics datasets may more accurately capture how a population of cells in a single tumor may respond to perturbations. Of course, at the same time, single cell phosphoproteomic measurements are not yet readily available. Another limitation to this work is the lack of experimental data to validate the model predictions, i.e., could a gene or protein knockout or pharmacological inhibition of a target protein prevent or promote the model-predicted transition?

Collecting time-course RNA sequencing and/or proteomic measurements from GBM cells either in the presence of drugs (inhibitors of proposed cell state drivers) or with proteins knocked out, would serve as an appropriate next step. Indeed, some of the proteins our model has predicted to be critical for cell state transitions have been previously identified as critical for certain phenotypes. For example, *STAT3* activity has been associated with the mesenchymal phenotype [61].

In conclusion, we built a Boolean model from a multi-omics dataset which permitted the definition of the four Suva cell states, previously identified in GBM patients. The ensuing model simulations supported by machine learning analyses of an independent GBM patient dataset provided hypotheses of the key drivers of cell state transitions and lay the groundwork for future experimental validation *in vitro* as well as the expanding the scope of the model to include the effect of other cell types in the microenvironment.

## METHODS

### Data Sources

Publicly available snRNA-seq, bulk RNA-seq, and phosphoproteomic data from healthy brain and GBM tissue were downloaded from the CPTAC portal (https://proteomics.cancer.gov/data-portal) as originally published in Wang et al. (2021) [34]. These data – referred to as the Wang dataset – were used in different analysis streams that culminated in a protein-protein interaction network (PPIN) for each of the four Neftel cell states or phenotypes. Each analysis stream is described in detail below. Bulk RNAseq and clinical data were downloaded from the Glioma Longitudinal Analysis (GLASS) Consortium (http://www.synapse.org/glass [62]. These data were used as a test set for machine learning algorithms.

### Classification of Patient-Specific Dominant Cell State

#### Single Cell Preprocessing

Eighteen snRNA samples (filtered feature matrices) from wild type (WT) IDH1 GBM patients as part of the Wang dataset, were imported into R via the Seurat package (v4) [35] for analysis. Cells in which <200 genes were identified and cells which express > 10,000 genes were excluded from the analysis. Any cell with mitochondrial gene expression exceeding 10% was excluded as these cells were likely apoptotic.

Genes that are expressed in <10 cells across all samples were filtered out due to low expression across the population. Cells were then split into immune and non-immune cells using CD45 (pan-leukocyte marker) expression as a biomarker. Non-immune cells were further analyzed in Seurat, while immune cells were excluded, as we are primarily concerned with cancer cell phenotype.

#### Data Transformation

The remaining feature counts were divided by the total feature counts in each cell and multiplied by a scale factor of 10,000. The result was transformed with the natural log +1.

#### Variable Features

The 3,000 most variable genes were identified with the variance stabilization transformation (VST). VST uses polynomial regression to fit a curve to the relationship of the log of the variance and mean of the dataset. The gene values were then standardized using the fitted line. Gene variance was estimated by using the standardized curve and any values exceeding the maximum (set to the square root of the number of cells) is set to this maximum.

The 3,000 most variable genes are then linear transformed in the following manner:

1. The average expression of each gene between all cells is 0
2. The variance of each gene between all cells is 1

#### Principal Component Analysis (PCA)

The 3,000 scaled genes are used to reduce the dimension of the dataset via principal component analysis (PCA), and the first 12 principal components were used.

#### Clustering and Labeling

Using the 12 principal components, a K-nearest neighbor (KNN) graph is built by calculating the Euclidean distance and Jaccard similarity. Cells are then clustered using the Louvain algorithm and a resolution of 0.5. 40 clusters were identified in this analysis. The Wilcoxon rank sum test was used to find differentially expressed genes amongst the clusters. Clusters were labeled using meta-modules as defined by Neftel et al. (2019) [28] and Wang et al (2021) [34]. A hypergeometric test was used to evaluate whether each meta-module was overrepresented in each cluster and clusters were labeled with the most overrepresented meta-module (-log(p-value).

### Deconvolution

The Bisque [36] package in R was used to deconvolve 74 bulk RNAseq samples from wtIDH1 patients and estimate cell type compositions. Read counts were normalized by counts-per-million (CPM) and relative abundances were calculated by averaging gene CPM within each cell type. A linear relationship was then defined between each gene’s single cell expression and bulk expression. Single cell expressions were taken from the eighteen patient samples previously analyzed. In this manner, cell type compositions can be estimated for both the eighteen patients with snRNA-seq data and the 74 patients with bulk RNAseq data thereby yielding a total of 92 patients.

### Patient Classification

Based on the proportional cell type composition estimates, each patient was classified into a dominate cell state–-one of four phenotypes as previously defined by Neftel et al [28]: Mesenchymal-like (MES-like), astrocyte-like (AC-like), Neural-progenitor-like (NPC-like), and Oligodendrocyte-progenitor-like (OPC-like).

### COSMOS PIPELINE

Causal Oriented Search of Multi-Omics Space (COSMOS) is a computational package utilizing multi-omics data to generate mechanistic hypotheses relevant to disease, including cancer [39]. The COSMOS pipeline outputs a directed PPIN as described below. The PPIN includes details on whether each protein node, including transcription factors, is up- or down–regulated and whether each interaction or edge between proteins is activating or inhibiting. Each of the four cell states was analyzed separately with this pipeline, utilizing Bulk RNAseq and phosphoproteins.

### Bulk RNAseq Analysis

RNAseq data were filtered for low expressed genes; any gene with read counts <50 was excluded from the data set. Genes were normalized using the median of ratios. *DESeq2* [38] was used to analyze remaining genes and a differential analysis was conducted between genes from healthy brain and genes from each of the four phenotypes. A Benjamini-Hochberg correction was applied and any gene with an adjusted p-value < 0.05 is considered differentially expressed.

### Phosphoproteomic Analysis

Phosphoproteins which were expressed in less than 25% of samples were excluded from the analysis. A differential analysis was performed using *limma* ([37]. A Benjamini-Hochberg correction was applied and any phosphoprotein with an adjusted p-value < 0.05 is differentially expressed.

### Footprint Based Activity Estimation

The *DoRothEA* R package [63] was used to download transcription factor-gene interactions. TF-gene relationships are scored from A-E based on the available evidence in literature, with A being the highest score. Interactions categorized with confidence levels of A-C were included in the analysis which includes all TF-gene relationships that are defined as high confidence (A), likely confidence (B), or medium confidence (C). High confidence interactions include interactions that are supported by four independent sources of evidence (curated databases, ChIP-seq experiments, transcription factor binding sites (TFBS), and genotype-tissue expression (GTEx). Likely confidence interactions include three of these lines of evidence, while medium confidence interactions include two. Protein-protein interactions were downloaded using *OmniPath* [64]. The WMEAN algorithm from the *decoupleR* R package [40] was used to calculate enrichment scores and activity scores for each transcription factor and protein.

### Prior Knowledge Network (PKN)

In concordance with COSMOS [39], the prior knowledge network was constructed from three databases: *STICHdb* (http://stitch.embl.de/), *OmniPath* [64], and ReCon3D [65]. *STICHdb* and *Recon3D* primarily contain interactions between metabolites and enzymes while *OmniPath* contains protein-protein interactions. Since we do not utilize metabolomic data in this study, most of the interactions in our networks come from *OmniPath*. The PKN includes a total of 76,638 interactions.

### COSMOS

The PKN and activity scores for both proteins and transcription factors are input into COSMOS. COSMOS uses CARNIVAL [66] to convert the data into an integer linear programming (ILP) problem and the IBM CPLEX ILP solver (https://www.ibm.com/products/ilog-cplex-optimization-studio/cplex-optimizer) was used to optimize the objective function. COSMOS was run in both forward and backward mode for each of the four phenotypes and the two runs were combined to generate the final PPI network for each of the four cell states.

### Boolean Network Construction

All four COSMOS networks were then combined into one final PPIN that was used to derive a Boolean logic model. The networks were combined by incorporating all unique interactions among the four networks. To elucidate the logical formalisms (e.g. AND vs OR gates) of the final network, MaBoSS (Version 2.0) [42] was used to conduct steady-state simulations. For each phenotype, input nodes (species in the network without any upstream effectors) were set to either “ON” (1) or “OFF” (0) depending on their state for each respective phenotype. Proteins which are upregulated according to COSMOS are set to “ON”, while proteins which are downregulated are set to “OFF”. The model was then run to steady-state. For a logical formalism to be considered correct, the steady-state of the simulation must fully match the desired phenotype as described by the COSMOS PPI networks. In other words, each individual node must be correctly in the “ON” or “OFF” state, as defined by our COSMOS analysis.

### Knockout Simulations

Knockout simulations were performed in MaBoSS utilizing the Boolean model. Starting at steady-state, each node which is in the “ON” state, is turned “OFF” one simulation at a time and the model is run until a new steady-state is reached. These simulations were performed from the steady-state of all four cell states or phenotypes resulting in 146 total simulations. Each simulation was evaluated for changes as compared to the baseline steady-state (Hamming Distance) for each phenotype and z-normalized. Each new steady-state resulting from the knockouts was classified into one of the four cell states by calculating the overall similarity between itself and the original COSMOS cell states.

### Validation of Model Simulations using Machine Learning

#### Model Training

Results from the knockout simulations were as the training data set to a machine learning model capable of predicting the predominant phenotype in a clinical cohort. The state of each node (protein) in the model was used as the input features to four different machine learning algorithms: multinomial regression, k-nearest neighbor (KNN), random forest (RF), and XGBoost. For multinomial regression, elastic net regularization was applied with an alpha = 0.14, which was optimized with a 10-fold cross validation on the training data [67]. For KNN, a 5-fold cross validation was used with recursive feature elimination to determine the optimal number of features (80) and neighbors (13). For both RF and XGBoost, the *Boruta* feature selection algorithm was used to select the most important features with a maximum runs of 100 [68]. The *Random Forest* package in R was used for the RF model with the following parameters changed from default: ntree (number of trees) = 1000 and mtry (number of features sampled at each tree split) = 35. XGBoost was implemented in Python with the following hyperparameters changed from default: n_estimators (number of trees) = 100, learning_rate = 0.1, max_depth (tree depth) = 1, and subsample (proportion of samples used per tree) = 0.1.

#### Model Testing and Interpretation

Bulk RNAseq from 201 wtIDH1 tumors from the GLASS Consortium were used to test model performance. These samples included 107 primary (untreated tumors) and 114 recurrent tumors. Many samples were from patients that had both primary and recurrent tumors. Using the same deconvolution method as previously described, each sample was classified as one of the four cell states. Next, log-fold changes for each sample were calculated as compared to healthy brain tissue. Footprint based activity measurements were estimated in the same manner as before using the *DoRothEA* R package. Since phosphoproteomic data was not available for the GLASS samples, we derived a simple ratio/relationship between phosphoproteomic activity and RNAseq activity estimates using the Wang dataset:

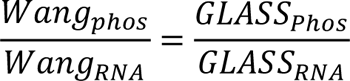

In this manner, we were able to obtain full activity estimates for the GLASS dataset. These activity estimates were then converted to Boolean values (positive activity = 1, negative activity = 0) to match the simulation data and to test the four machine learning models. The TreeSHAP method for calculating Shapley values from decision tree models was used with default parameters for model interpretation [69].

## Supporting information

Differentially Expressed Genes

Differentially Expressed Phosphoproteins

Protein-Protein Interaction Networks

## Author contributions

Conceptualization: B.B, J.M.G. Methodology: B.B., M.R.B., J.M.G. Investigation: B.B. Supervision: M.R.B, J.M.G., Writing-original draft: B.B., Writing-review and editing: B.B., M.R.B, J.M.G

## Acknowledgements

Brandon Bumbaca was funded, in part, with research support from the University at Buffalo School of Pharmacy and Pharmaceutical Sciences. MRB thanks NIH / NIGMS for funding (R35GM141891).

## Data availability

SnRNA-seq, bulk RNA-seq, and phosphoproteomic data originally published in Wang et al. (2021) [34], are available at the CPTAC portal (https://proteomics.cancer.gov/data-portal). Bulk RNAseq and clinical data from the Glioma Longitudinal Analysis (GLASS) Consortium are available online (http://www.synapse.org/glass) [62]. Differentially expressed genes and phosphoproteins from each cell state are available in the supplementary material. Details of all the PPINs are also available in the supplementary material and any other relevant data is available upon request.

## Code availability

All code required to generate simulations and figures are available upon request.

## Competing interests

All authors declare no financial or non-financial competing interests.

## Notes

### Competing Interest Statement

The authors have declared no competing interest.

