## Supplemental_Information for "Network Analyses of Brain Tumor Patients’ Multiomic Data Reveals Pharmacological Opportunities to Alter Cell State Transitions"

### SUPPLEMENTARY INFORMATION

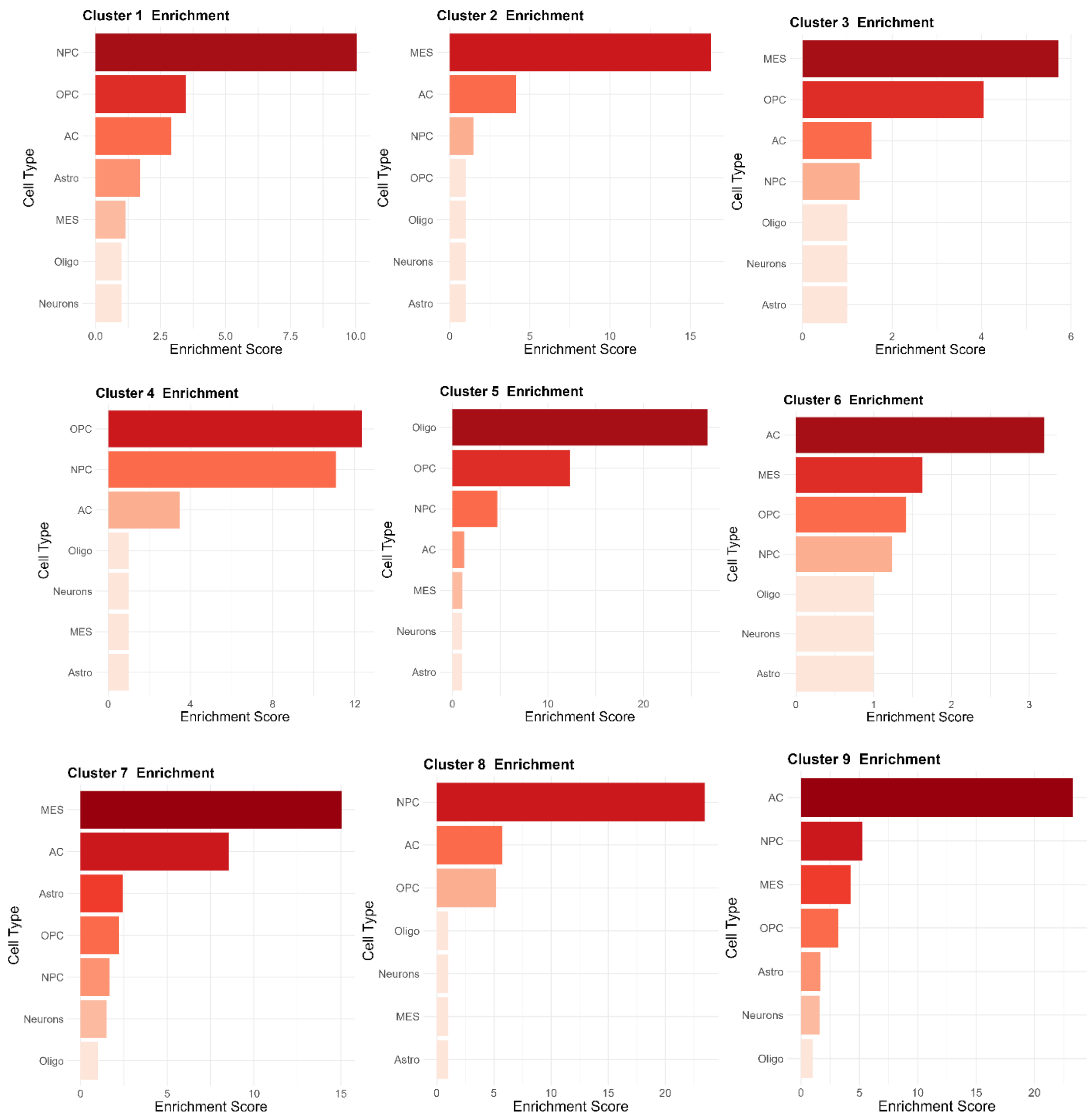

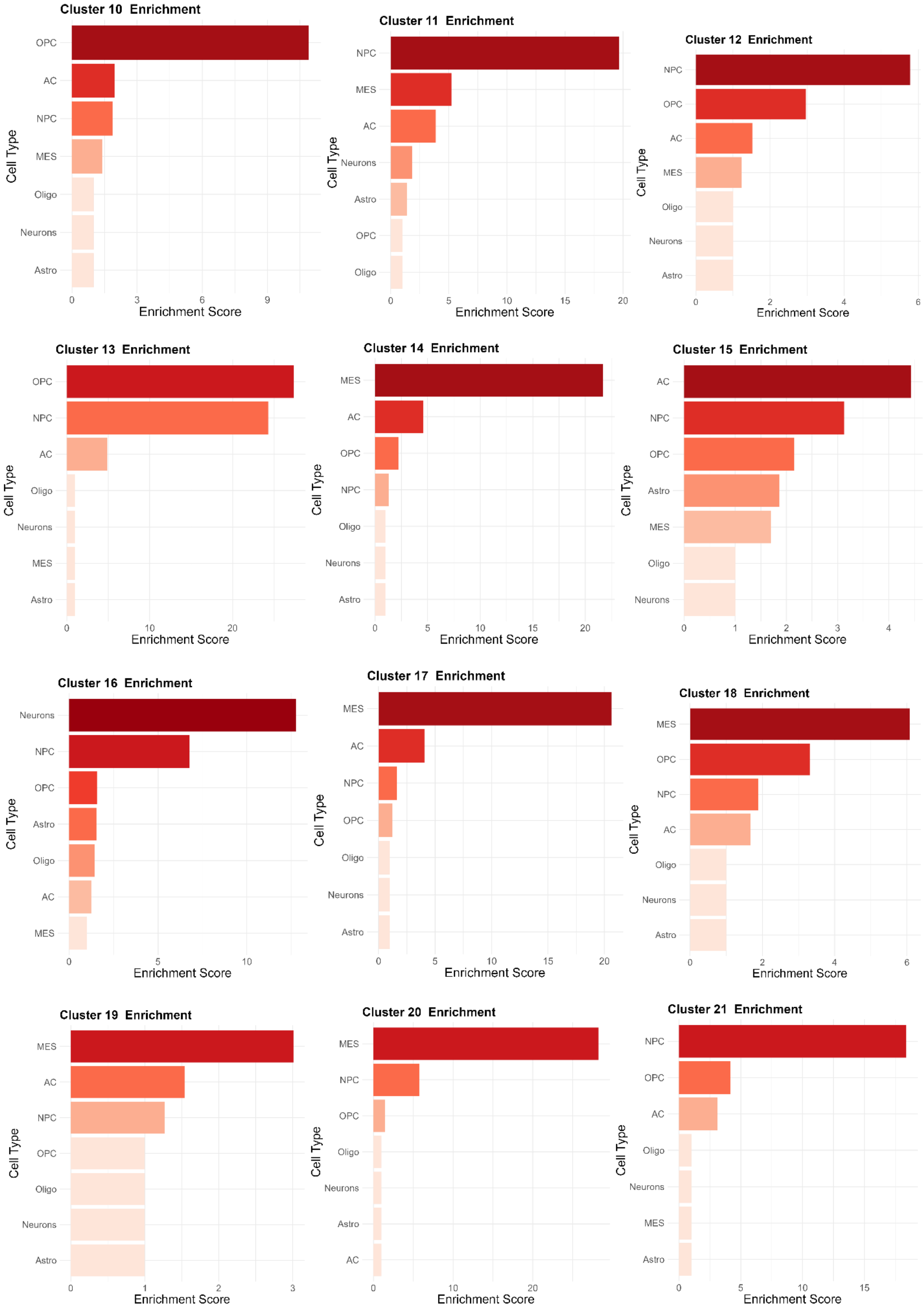

Cluster 10 Enrichment
OPC
AC
NPC
MES
Oligo
Neurons
Astro
Cell Type
0 3 6 9
Enrichment Score
Cluster 11 Enrichment
NPC
MES
AC
Neurons
Astro
OPC
Oligo
Cell Type
0 5 10 15 20
Enrichment Score
Cluster 12 Enrichment
NPC
OPC
AC
MES
Oligo
Neurons
Astro
Cell Type
0 2 4 6
Enrichment Score
Cluster 13 Enrichment
OPC
NPC
AC
Oligo
Neurons
MES
Astro
Cell Type
0 10 20
Enrichment Score
Cluster 14 Enrichment
MES
AC
OPC
NPC
Oligo
Neurons
Astro
Cell Type
0 5 10 15 20
Enrichment Score
Cluster 15 Enrichment
AC
NPC
OPC
Astro
MES
Oligo
Neurons
Cell Type
0 1 2 3 4
Enrichment Score
Cluster 16 Enrichment
Neurons
NPC
OPC
Astro
Oligo
AC
MES
Cell Type
0 5 10
Enrichment Score
Cluster 17 Enrichment
MES
AC
NPC
OPC
Oligo
Neurons
Astro
Cell Type
0 5 10 15 20
Enrichment Score
Cluster 18 Enrichment
MES
OPC
NPC
AC
Oligo
Neurons
Astro
Cell Type
0 2 4 6
Enrichment Score
Cluster 19 Enrichment
MES
AC
NPC
OPC
Oligo
Neurons
Astro
Cell Type
0 1 2 3
Enrichment Score
Cluster 20 Enrichment
MES
NPC
OPC
Oligo
Neurons
Astro
AC
Cell Type
0 10 20
Enrichment Score
Cluster 21 Enrichment
NPC
OPC
AC
Oligo
Neurons
MES
Astro
Cell Type
0 5 10 15
Enrichment Score

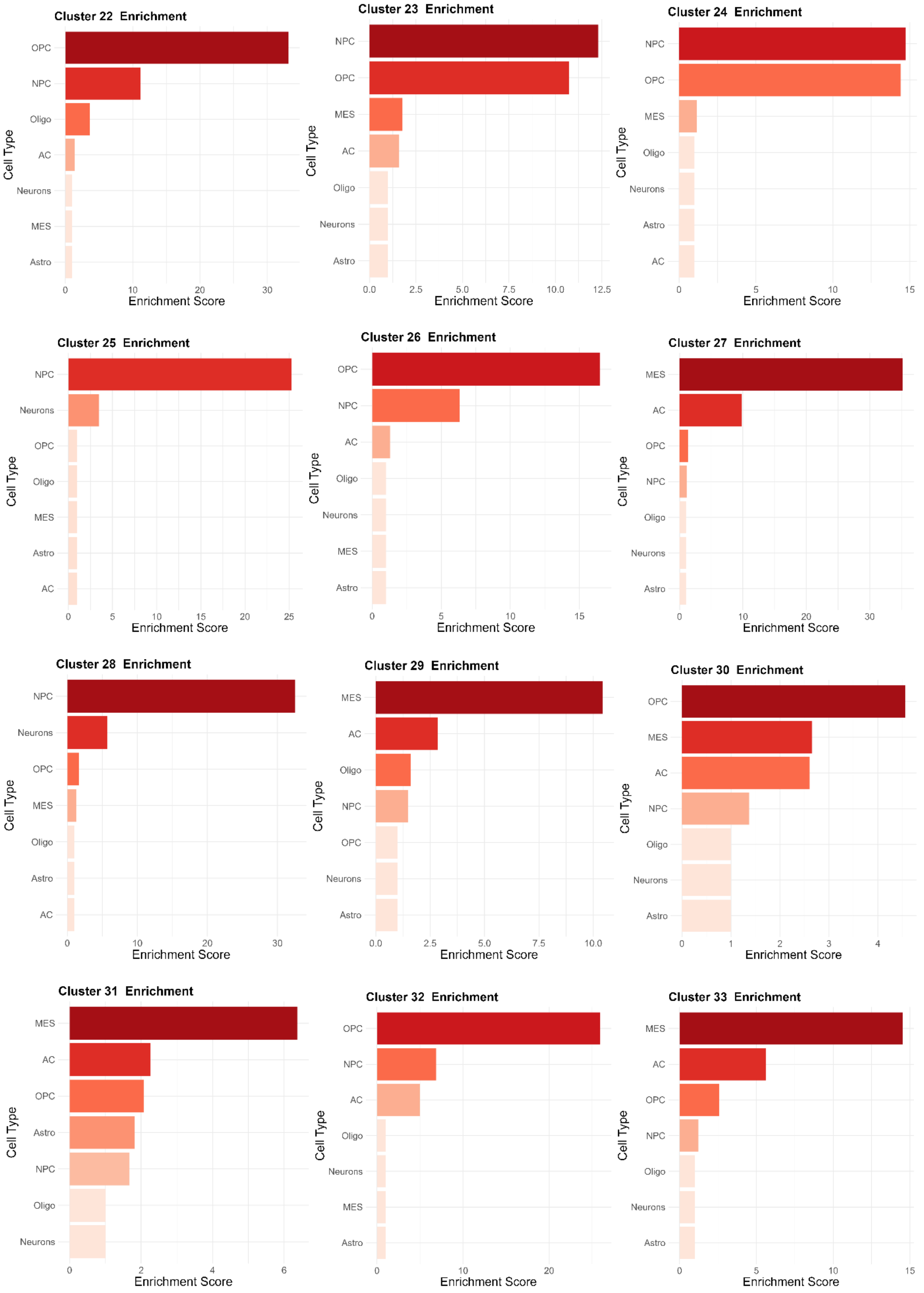
Cluster 22 Enrichment
OPC
NPC
Oligo
AC
Neurons
MES
Astro
Cell Type
0
10
20
30
Enrichment Score
Cluster 23 Enrichment
NPC
OPC
MES
AC
Oligo
Neurons
Astro
Cell Type
0.0
2.5
5.0
7.5
10.0
12.5
Enrichment Score
Cluster 24 Enrichment
NPC
OPC
MES
Oligo
Neurons
Astro
AC
Cell Type
0
5
10
15
Enrichment Score
Cluster 25 Enrichment
NPC
Neurons
OPC
Oligo
MES
Astro
AC
Cell Type
0
5
10
15
20
25
Enrichment Score
Cluster 26 Enrichment
OPC
NPC
AC
Oligo
Neurons
MES
Astro
Cell Type
0
5
10
15
Enrichment Score
Cluster 27 Enrichment
MES
AC
OPC
NPC
Oligo
Neurons
Astro
Cell Type
0
10
20
30
Enrichment Score
Cluster 28 Enrichment
NPC
Neurons
OPC
MES
Oligo
Astro
AC
Cell Type
0
10
20
30
Enrichment Score
Cluster 29 Enrichment
MES
AC
Oligo
NPC
OPC
Neurons
Astro
Cell Type
0.0
2.5
5.0
7.5
10.0
Enrichment Score
Cluster 30 Enrichment
OPC
MES
AC
NPC
Oligo
Neurons
Astro
Cell Type
0
1
2
3
4
Enrichment Score
Cluster 31 Enrichment
MES
AC
OPC
Astro
NPC
Oligo
Neurons
Cell Type
0
2
4
6
Enrichment Score
Cluster 32 Enrichment
OPC
NPC
AC
Oligo
Neurons
MES
Astro
Cell Type
0
10
20
Enrichment Score
Cluster 33 Enrichment
MES
AC
OPC
NPC
Oligo
Neurons
Astro
Cell Type
0
5
10
15
Enrichment Score

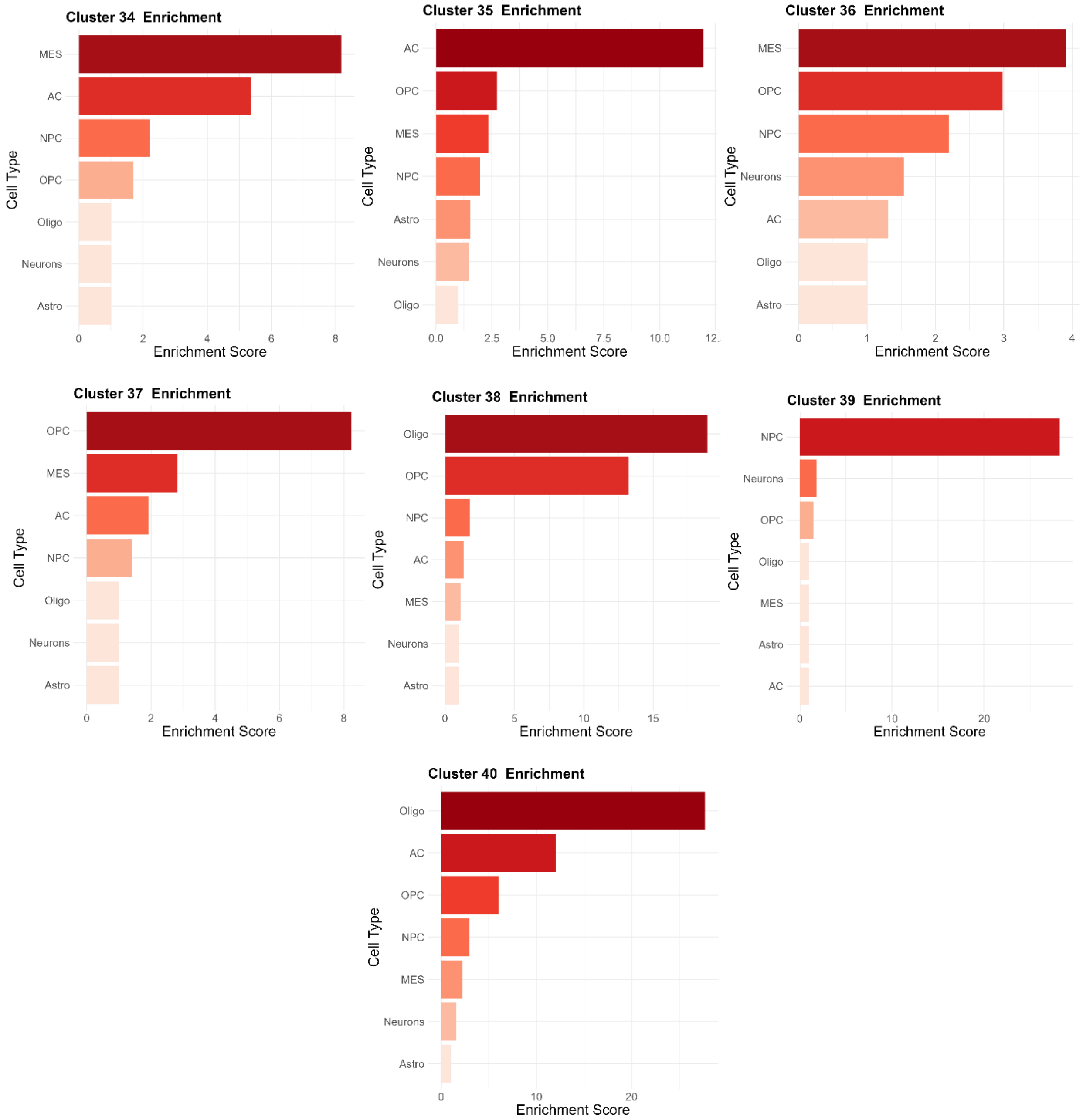


**Supplementary Figure 1 (S1). Single Cell Cluster Enrichment.** Clusters were labeled using meta-modules as defined by Neftel et al. (2019) [1] and Wang et al (2021) [2]. A hypergeometric test was used to evaluate whether each meta-module was overrepresented in each cluster and clusters were labeled with the most overrepresented meta-module (-log(p-value).

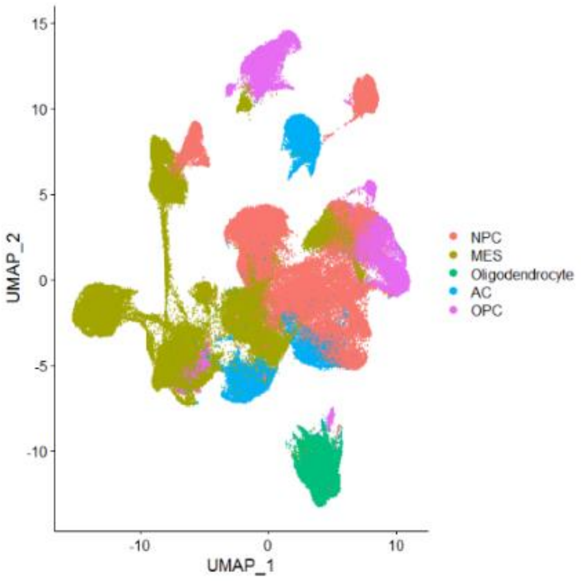


**Supplementary Figure 2 (S2). UMAP of snRNA-seq Data.**

| **Species** | **Logic** |
|---|---|
| ABL1 | = (!ATM & !CDK1 & !CDK2 & !FYN & !LCK & !PAK2 & !PRKCA & !PRKDC & !RFX1 & SRC) \| (!ATM & !CDK1 & !CDK2 & !FYN & !LCK & !PAK2 & !PRKCA & !PRKDC & RFX1) \| (!ATM & !CDK1 & !CDK2 & !FYN & !LCK & !PAK2 & !PRKCA & PRKDC) \| (!ATM & !CDK1 & !CDK2 & !FYN & !LCK & !PAK2 & PRKCA) \| (!ATM & !CDK1 & !CDK2 & !FYN & !LCK & PAK2) \| (!ATM & !CDK1 & !CDK2 & !FYN & LCK) \| (!ATM & !CDK1 & !CDK2 & FYN) \| (!ATM & !CDK1 & CDK2) \| (!ATM & CDK1) \| (ATM) |
| ATR | ABL1 |
| EGFR | (ABL1 \| AR \| EGF \| FGF2 \| FYN \| HGF \| IRF1 \| JUN \| JUNB \| NFKB1 \| RARA \| RELA) |
| AGT | (!FOS & HIF1A) \| (FOS) |
| AHR | AHR |
| AKT1 | (AR \| ATM \| AURKA \| FGF2 \| LCK \| PRKDC \| RAC1 \| SRC \| TBK1 \| TNF) & (!TP53 \| !PPP2CA) |
| AKT2 | (SRC \| INSR \| PRKDC \| TBK1) & (!PPP2CA \| !PTEN) |
| AKT3 | (!MAP3K7 & !PPP2CA & !PRKCZ & !PRKDC & !PTEN & !PTK2B & !SRC & TBK1) \| (!MAP3K7 & !PPP2CA & !PRKCZ & !PRKDC & !PTEN |

| | |
|---|---|
| | & !PTK2B & SRC) \| (!MAP3K7 & !PPP2CA & !PRKCZ & !PRKDC & !PTEN & PTK2B) \| (!MAP3K7 & !PPP2CA & !PRKCZ & !PRKDC & PTEN) \| (!MAP3K7 & !PPP2CA & !PRKCZ & PRKDC) \| (!MAP3K7 & !PPP2CA & PRKCZ) \| (MAP3K7 & !PPP2CA) |
| AR | AR |
| ARAF | PAK1 |
| ARNT | ARNT |
| ATM | (!ATR & !CDK5 & !E2F1 & !EGF & !FOXO3 & !PPP2CA & !RPS6KA2 & TNF) \| (!ATR & !CDK5 & !E2F1 & !EGF & !FOXO3 & !PPP2CA & RPS6KA2) \| (!ATR & !CDK5 & !E2F1 & !EGF & FOXO3 & !PPP2CA) \| (!ATR & !CDK5 & !E2F1 & EGF & !PPP2CA) \| (!ATR & !CDK5 & E2F1 & !PPP2CA) \| (!ATR & CDK5 & !PPP2CA) \| (ATR & !PPP2CA) |
| AURKA | (!E2F1 & PRKACA) \| (E2F1) |
| BCL6 | BCL6 |
| BDNF | TNF |
| BRAF | (!AKT1 & !AKT2 & !AKT3 & !PAK4 & PPP2CA) \| (!AKT1 & !AKT2 & !AKT3 & PAK4) |
| BUB1 | CDK1 |
| CAMK2B | CAMKK1 |
| CAMKK1 | (!CDK6 & CDK5) \| (CDK6) |
| CDK1 | (E2F1 & E2F4) & (FOS \| SP1 \|!TP53 \| !GSK3B \| !PIN1 \| !MAPK1) |
| CDK2 | (!AKT1 & !AKT3 & !MAPK3 & MITF & !TGFB1) \| (!AKT1 & !AKT3 & MAPK3 & !TGFB1) \| (!AKT1 & AKT3 & !TGFB1) \| (AKT1 & !TGFB1) |
| CDK4 | (!FOS & !GSK3B & !JUN & !MAX & !MYC & !NFKB1 & !RELA & STAT3 & !YES1) \| (!FOS & !GSK3B & !JUN & !MAX & !MYC & !NFKB1 & RELA & !YES1) \| (!FOS & !GSK3B & !JUN & !MAX & !MYC & NFKB1 & !YES1) \| (!FOS & !GSK3B & !JUN & !MAX & MYC & !YES1) \| (!FOS & !GSK3B & !JUN & MAX & !YES1) \| (!FOS & !GSK3B & JUN & !YES1) \| (FOS & !GSK3B & !YES1 |
| CDK6 | (!GSK3B & !MAX & !MYC & RELA & !SP1) \| (!GSK3B & !MAX & MYC & !SP1) \| (!GSK3B & MAX & !SP1) |
| CDK5 | FYN |
| CDK9 | AKT1 & MAPK3 |
| CEBPA | CEBPA |

| | |
|---|---|
| CHEK1 | (AKT1 \| ATM \| ATR \| CDK1 \| E2F1 \| CDK2 \| MAP3K8) & (!BCL6 \| !PPP2CA \| !TP53) |
| CHEK2 | CDK9 & PPP2CA |
| CHUK | (!AKT1 & !AKT2 & !AKT3 & !MAPK3 & !NR2C2 & !PRKCB & !PRKCE & SRC & !TP53) \| (!AKT1 & !AKT2 & !AKT3 & !MAPK3 & !NR2C2 & !PRKCB & PRKCE & !TP53) \| (!AKT1 & !AKT2 & !AKT3 & !MAPK3 & !NR2C2 & PRKCB & !TP53) \| (!AKT1 & !AKT2 & !AKT3 & !MAPK3 & NR2C2 & !TP53) \| (!AKT1 & !AKT2 & !AKT3 & MAPK3 & !TP53) \| (!AKT1 & !AKT2 & AKT3 & !TP53) \| (!AKT1 & AKT2 & !TP53) \| (AKT1 & !TP53) |
| CREB1 | !ATM |
| CREBBP | (!NFKB1 & !RUNX2 & !SMAD3 & SMAD4 & !YY1) \| (!NFKB1 & !RUNX2 & SMAD3 & !YY1) \| (!NFKB1 & RUNX2 & !YY1) \| (NFKB1 & !YY1) |
| CSK | (!CREBBP & !PRKAR2A & !PRKAR2B & VEGFA) \| (!CREBBP & !PRKAR2A & PRKAR2B) \| (!CREBBP & PRKAR2A) \| (CREBBP) |
| CSNK1E | PLK1 |
| CSKN1A1 | (!PPP3CA & PPP3CB) \| (PPP3CA) |
| CSNK2A1 | (MAPK14 & !NOLC1) |
| DAPK1 | !RPS6KA3 |
| DDX58 | !CSNK2A1 |
| DYRK2 | ATM |
| E2F1 | ATM |
| EGF | (!FYN & LCK) \| (FYN) |
| E2F4 | E2F4 |
| EGR1 | EGR1 |
| ELK1 | ELK1 |
| ETV1 | ETV1 |
| FGF2 | (HIF1A \| STAT1 \| TNF) & (!TP53 \| !SRC) |
| FOS | FOS |
| FOXO1 | FOXO1 |
| FOXO3 | FOXO3 |
| FYN | CSK |
| GRK5 | CDK1 |
| GSK3A | PRKCZ |
| GSK3B | (!AKT1 & !AKT2 & !AKT3 & MAPK1 & !PRKACA & !PRKCA & !PRKCZ & !RPS6KA5) |
| HGF | OSM & !PTEN |
| IGF1R | !TP53 |
| HIF1A | ATM |

| | |
|---|---|
| IKBKB | (!AKT1 & !AKT2 & !AKT3 & !NR2C2 & !PLK1 & !PRKCA & !PRKCB & !PRKCE & !PRKCZ & !SRC & TBK1) \| (!AKT1 & !AKT2 & !AKT3 & !NR2C2 & !PLK1 & !PRKCA & !PRKCB & !PRKCE & !PRKCZ & SRC) \| (!AKT1 & !AKT2 & !AKT3 & !NR2C2 & !PLK1 & !PRKCA & !PRKCB & !PRKCE & PRKCZ) \| (!AKT1 & !AKT2 & !AKT3 & !NR2C2 & !PLK1 & !PRKCA & !PRKCB & PRKCE) \| (!AKT1 & !AKT2 & !AKT3 & !NR2C2 & !PLK1 & !PRKCA & PRKCB) \| (!AKT1 & !AKT2 & !AKT3 & !NR2C2 & !PLK1 & PRKCA) \| (!AKT1 & !AKT2 & !AKT3 & NR2C2 & !PLK1) \| (!AKT1 & !AKT2 & AKT3 & !PLK1) \| (!AKT1 & AKT2 & !PLK1) \| (AKT1 & !PLK1) |
| IL1B | (!FOS & !JUN & !NFKB1 & !RELA & !STAT1 & STAT3) \| (!FOS & !JUN & !NFKB1 & !RELA & STAT1) \| (!FOS & !JUN & !NFKB1 & RELA) \| (!FOS & !JUN & NFKB1) \| (!FOS & JUN) \| (FOS) |
| INSR | (!CEBPA & PRKCE) \| (CEBPA) |
| IRF1 | IRF1 |
| JUNB | JUNB |
| JUN | JUN |
| LCK | FYN & MAPK3) \| (!CSK) |
| MAFB | MAFB |
| MAP2K1 | (EGFR \| MAP3K7 \| PAK2 \| RAC1) & (!CDK1 \| !PPP2CA) |
| MAP3K7 | (!PPP2CA) \| (TRAF6) |
| MAP3K8 | AKT1 |
| MAPK1 | (FGF2 \| YES1 \| IL1B \| PRKCA \| !SGK1) & (!HGF) |
| MAPK10 | !CDK5 & TRAF6 |
| MAPK12 | (LCK & EGF & CDK5) \| (PDPK1 & PGR) |
| MAPK11 | PAK1 & TRAF6 |
| MAPK13 | PGR |
| MAPK14 | MAP3K7 & MAPK11 & TRAF6 |
| MAPK8 | (PRKAA1 \| PRKAA2 \| PRKDC \| PRKCZ) & (MAP2K1 & MAP3K7) & (HGF \| TNF \| BDNF) & (RAC1 \| TGFBR2 \| PTK2B) |
| MAPK3 | (MAPK1 & (AHR \| FGF2 \| ETV1 \| MAFB) & (!WT1)) |
| MAPKAPK2 | (!MAPK12 & TNF) \| (MAPK12) |
| MAX | MAX |
| MITF | MITF |
| MYC | MYC |
| NFATC1 | NFACT1 |

| | |
|---|---|
| NFKB1 | NFKB1 |
| NOLC1 | RELA |
| NR2C2 | NR2C2 |
| OSM | STAT3 |
| PAK1 | PDPK1 |
| PAK2 | (!PRKCE & !SRC & TGFB1) \| (!PRKCE & SRC) \| (PRKCE) |
| PAK4 | (!PRKCE & YES1) \| (PRKCE) |
| PDPK1 | PTEN & IGF1R |
| PGR | PGR |
| PIM1 | (!PRKACA & !SP1 & !STAT1 & !STAT2 & STAT3) \| (!PRKACA & !SP1 & !STAT1 & STAT2) \| (!PRKACA & !SP1 & STAT1) \| (!PRKACA & SP1) \| (PRKACA) |
| PIM2 | (!PRKACA & STAT3) \| (PRKACA) |
| PIM3 | STAT5A |
| PIN1 | !DAPK1 |
| PLK1 | !ATM |
| PPARG | PPARG |
| PPP2CA | (!CREB1 & !SRC & TGFBR2) \| (CREB1 & !SRC) |
| PPP3CA | CAMK2B |
| PPP3CB | CAMK2B |
| PRKAA1 | (!CAMK2B & !CAMKK1 & MAP3K7) \| (!CAMK2B & CAMKK1) \| (CAMK2B) |
| PRKAA2 | MAP3K7 |
| PRKACA | (!AKT3 & !PPP2CA & RPS6KA5) \| (AKT3 & !PPP2CA) |
| PRKAR2A | CDK1 |
| PRKAR2B | CDK1 |
| PRKCA | IL1B & !PPP2CA |
| PRKCB | LCK |
| PRKCE | FYN & !SP1 & !STAT1 |
| PRKCZ | !PPP2CA |
| PRKD1 | PDPK1 |
| PRKDC | (!ATR & !EGFR & ATM) \| (!ATR & EGFR) \| (ATR) |
| PSEN1 | (!CDK5 & !ELK1 & PRKCB) \| (CDK5 & !ELK1) |
| PTEN | (TFAP2A \| (TP53 & LCK) \| (TP53 & PRKCA) \| (PRKCA & LCK)) & (!FOS \| !CSNK2A1) |
| PTK2B | (!CSK & !SRC & VEGFA) \| (!CSK & SRC) \| (CSK) |
| RAC1 | (!ABL1 & !EGFR & !FYN & !HGF & PTK2B & !TP53) \| (!ABL1 & !EGFR & !FYN & HGF & !TP53) \| (!ABL1 & !EGFR & FYN & !TP53) \| (!ABL1 & EGFR & !TP53) \| (ABL1 & !TP53) |
| RARA | RARA |

| | |
|---|---|
| RFX1 | RFX1 |
| RELA | RELA |
| RPS6KA2 | (!MAPK3 & PIM2 & RPS6KA4) \| (MAPK3 & RPS6KA4) |
| RPS6KA3 | (!FOXO1 & FYN) \| (FOXO1); |
| RPS6KA4 | (!MAPK12 & MAPK3) \| (MAPK12) |
| RPS6KA5 | (!MAP2K1 & MAPK1) \| (MAP2K1) |
| RUNX2 | RUNX2 |
| SGK1 | SRC |
| SMAD3 | SMAD3 |
| SMAD4 | SMAD4 |
| SP1 | !ATM |
| SRC | (!CAMK2B & !CDK5 & !CSK & !EGF & !PRKCB & !PRKCE & VEGFA) \| (!CAMK2B & !CDK5 & !CSK & !EGF & !PRKCB & PRKCE) \| (!CAMK2B & !CDK5 & !CSK & !EGF & PRKCB) \| (!CAMK2B & !CDK5 & !CSK & EGF) \| (!CAMK2B & !CDK5 & CSK) \| (!CAMK2B & CDK5) \| (CAMK2B) |
| STAT1 | STAT1 |
| STAT2 | STAT2 |
| STAT3 | STAT3 |
| STAT5A | MAPK8 |
| TBK1 | DDX58 |
| TFAP2A | TFAP2A |
| TFRC | (!ARNT & !HIF1A & MAX) \| (!ARNT & HIF1A) \| (ARNT) |
| TGFB1 | (!EGR1 & !FOS & !HIF1A & !JUN & !NFKB1 & !RELA & USF1 & !YY1) \| (!EGR1 & !FOS & !HIF1A & !JUN & !NFKB1 & RELA & !YY1) \| (!EGR1 & !FOS & !HIF1A & !JUN & NFKB1 & !YY1) \| (!EGR1 & !FOS & !HIF1A & JUN & !YY1) \| (!EGR1 & !FOS & HIF1A & !YY1) \| (!EGR1 & FOS & !YY1) \| (EGR1 & !YY1) |
| TGFBR2 | PDPK1 |
| TNF | (!CREBBP & !FOS & !JUN & !NFATC1 & !NFKB1 & !PTEN & RELA & !SP1) \| (!CREBBP & !FOS & !JUN & !NFATC1 & NFKB1 & !PTEN & !SP1) \| (!CREBBP & !FOS & !JUN & NFATC1 & !PTEN & !SP1) \| (!CREBBP & !FOS & JUN & !PTEN & !SP1) \| (!CREBBP & FOS & !PTEN & !SP1) \| (CREBBP & !PTEN & !SP1) |
| TP53 | TP53 |
| TRAF6 | !PSEN1 |

| | |
|---|---|
| USF1 | USF1 |
| VEGFA | (AR \| EGR1 \| ESR1 \| FOS \| HIF1A \| JUN \| MYC \| NFKB1 \| RELA \| SMAD3 \| SP1 \| STAT1 \| STAT3 \| TFAP2A) |
| WT1 | WT1 |
| YES1 | !CSK |
| YY1 | YY1 |

**Supplementary Figure 3 (S3). Boolean Logic Rules.** Logical rules that govern the Boolean Model. Symbol Meanings: **&** (AND Gate), **|** (OR Gate), **!** (NOT).

**a)**

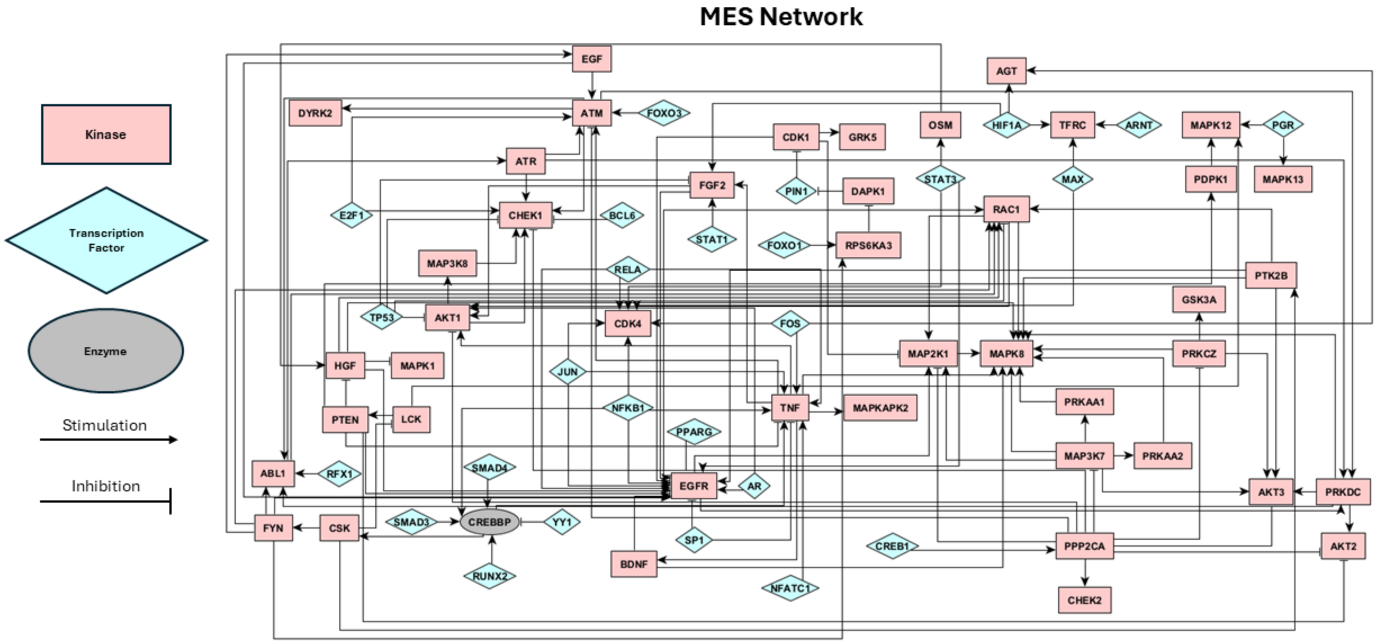

b)

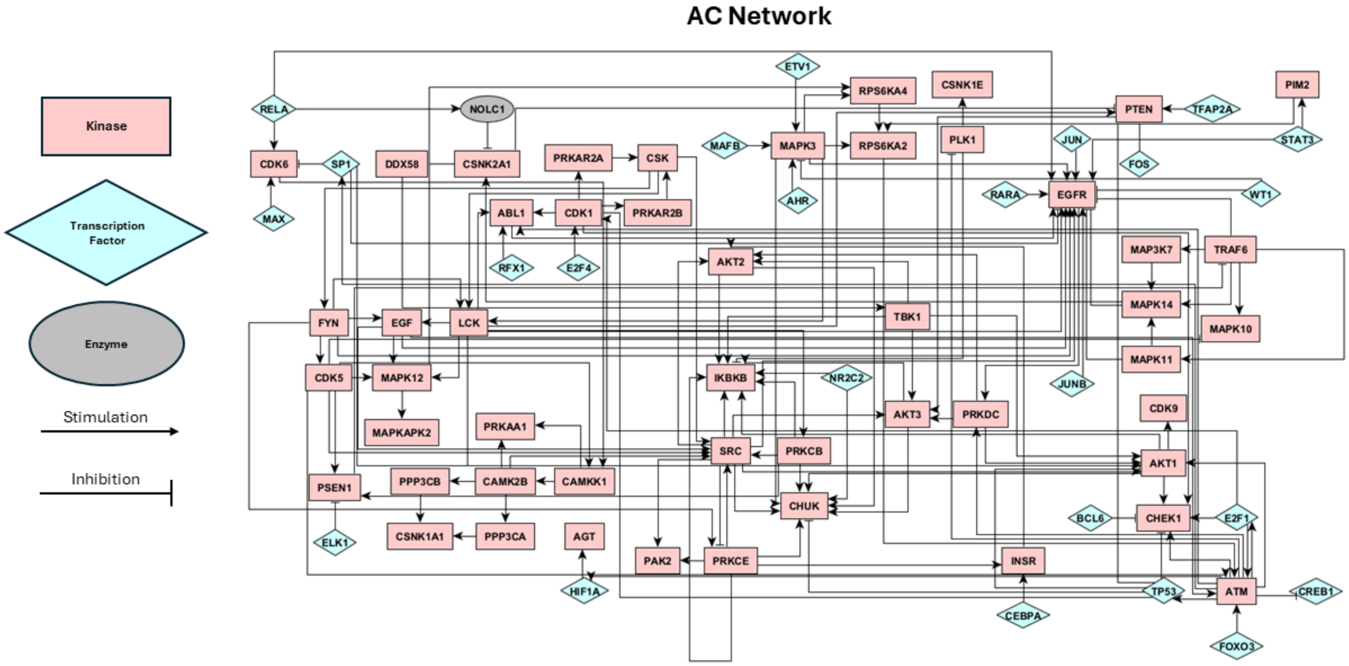
AC Network
Kinase
Transcription Factor
Enzyme
Stimulation
Inhibition
ETV1
CSNK1E
PIM2
RELA
NOLC1
RPS6KA4
PTEN
TFAP2A
CDK6
SP1
DDX58
CSNK2A1
PRKAR2A
CSK
MAFB
MAPK3
RPS6KA2
PLK1
JUN
STAT3
FOS
MAX
ABL1
CDK1
PRKAR2B
AHR
RARA
EGFR
WT1
RFX1
E2F4
AKT2
MAP3K7
TRAF6
TBK1
MAPK14
FYN
EGF
LCK
MAPK10
CDK5
MAPK12
IKBKB
NR2C2
MAPK11
JUNB
AKT3
PRKDC
CDK9
MAPKAPK2
PRKAA1
SRC
PRKCB
AKT1
PSEN1
PPP3CB
CAMK2B
CAMKK1
CHUK
BCL6
CHEK1
E2F1
ELK1
CSNK1A1
PPP3CA
AGT
PAK2
PRKCE
INSR
HIF1A
TP53
ATM
CREB1
CEBPA
FOXO3

c)

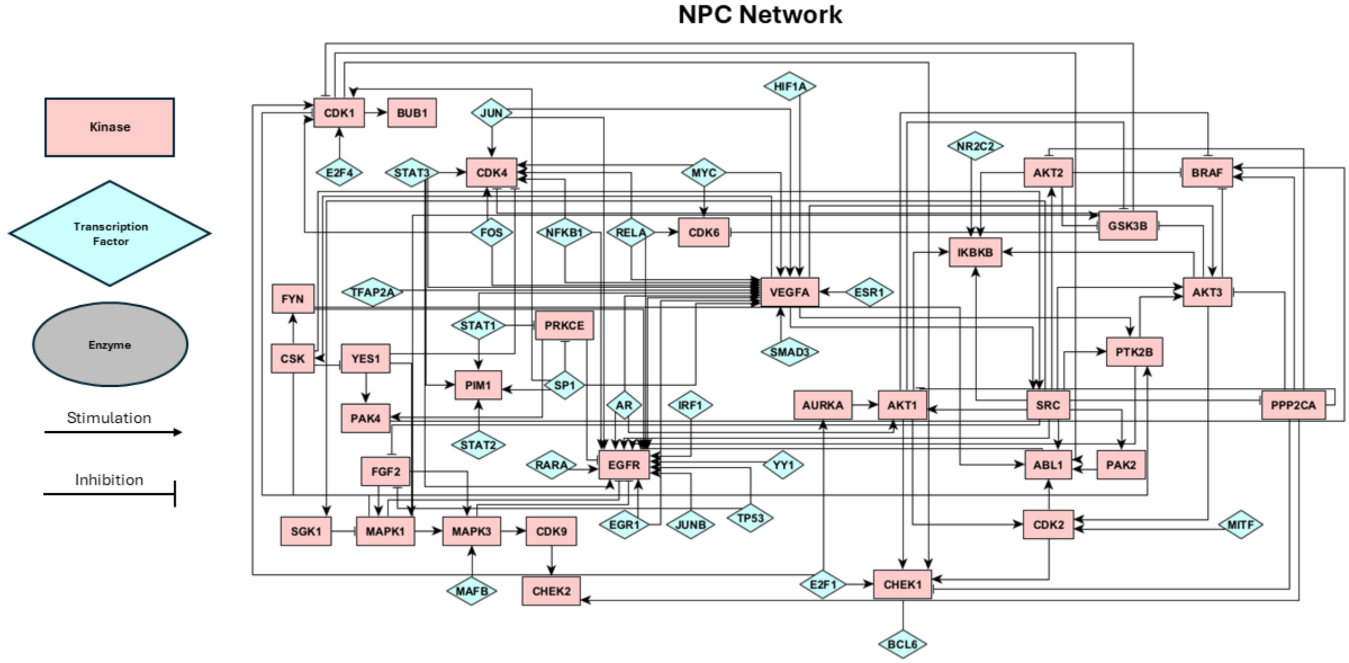
NPC Network
Kinase
Transcription Factor
Enzyme
Stimulation
Inhibition
HIF1A
CDK1
BUB1
JUN
NR2C2
E2F4
STAT3
CDK4
MYC
AKT2
BRAF
FOS
NFKB1
RELA
CDK6
GSK3B
IKBKB
TFAP2A
VEGFA
ESR1
AKT3
FYN
STAT1
PRKCE
PTK2B
CSK
YES1
SMAD3
PIM1
SP1
PAK4
AR
IRF1
AURKA
AKT1
SRC
PPP2CA
STAT2
RARA
EGFR
YY1
ABL1
PAK2
FGF2
EGR1
JUNB
TP53
SGK1
MAPK1
MAPK3
CDK9
CDK2
MITF
MAFB
CHEK2
E2F1
CHEK1
BCL6

d)

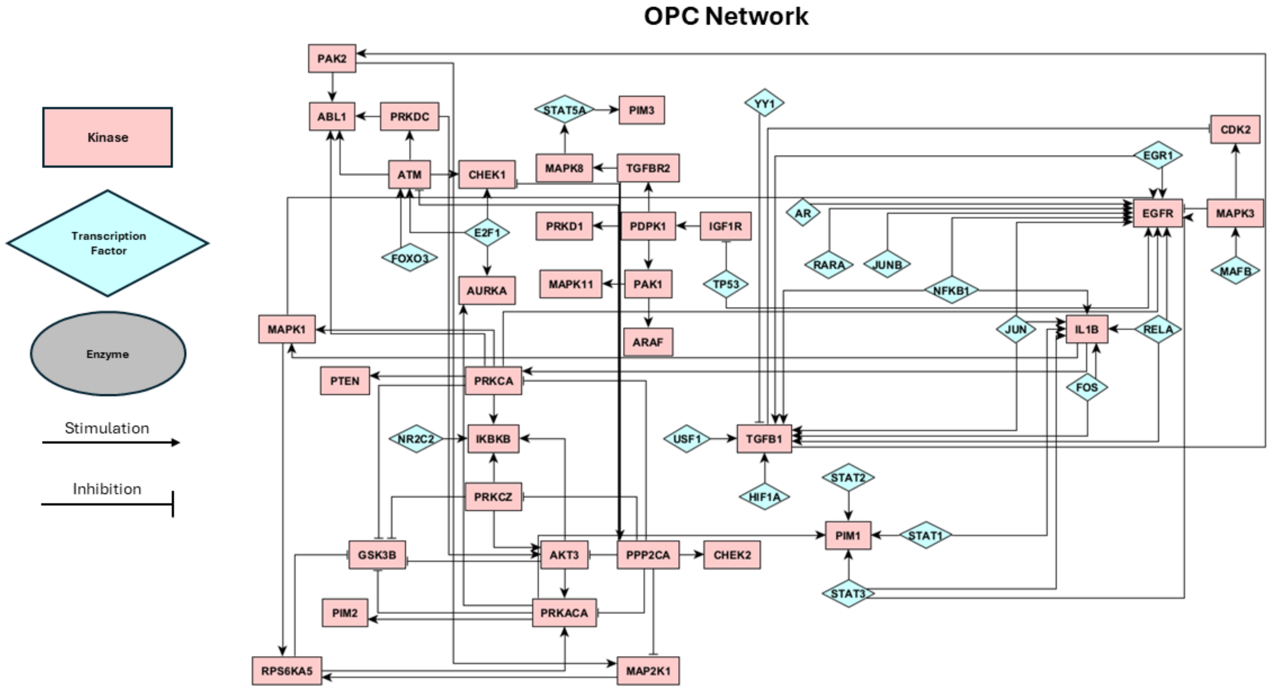
OPC Network
Kinase
Transcription Factor
Enzyme
Stimulation
Inhibition
PAK2
ABL1
PRKDC
STAT5A
PIM3
YY1
CDK2
ATM
CHEK1
MAPK8
TGFBR2
EGR1
AR
EGFR
MAPK3
PRKD1
PDPK1
IGF1R
E2F1
FOXO3
RARA
JUNB
MAFB
AURKA
MAPK11
PAK1
TP53
NFKB1
MAPK1
ARAF
JUN
IL1B
RELA
PTEN
PRKCA
FOS
NR2C2
IKBKB
USF1
TGFB1
STAT2
PRKCZ
HIF1A
GSK3B
AKT3
PPP2CA
CHEK2
PIM1
STAT1
PIM2
PRKACA
STAT3
RPS6KA5
MAP2K1

e)

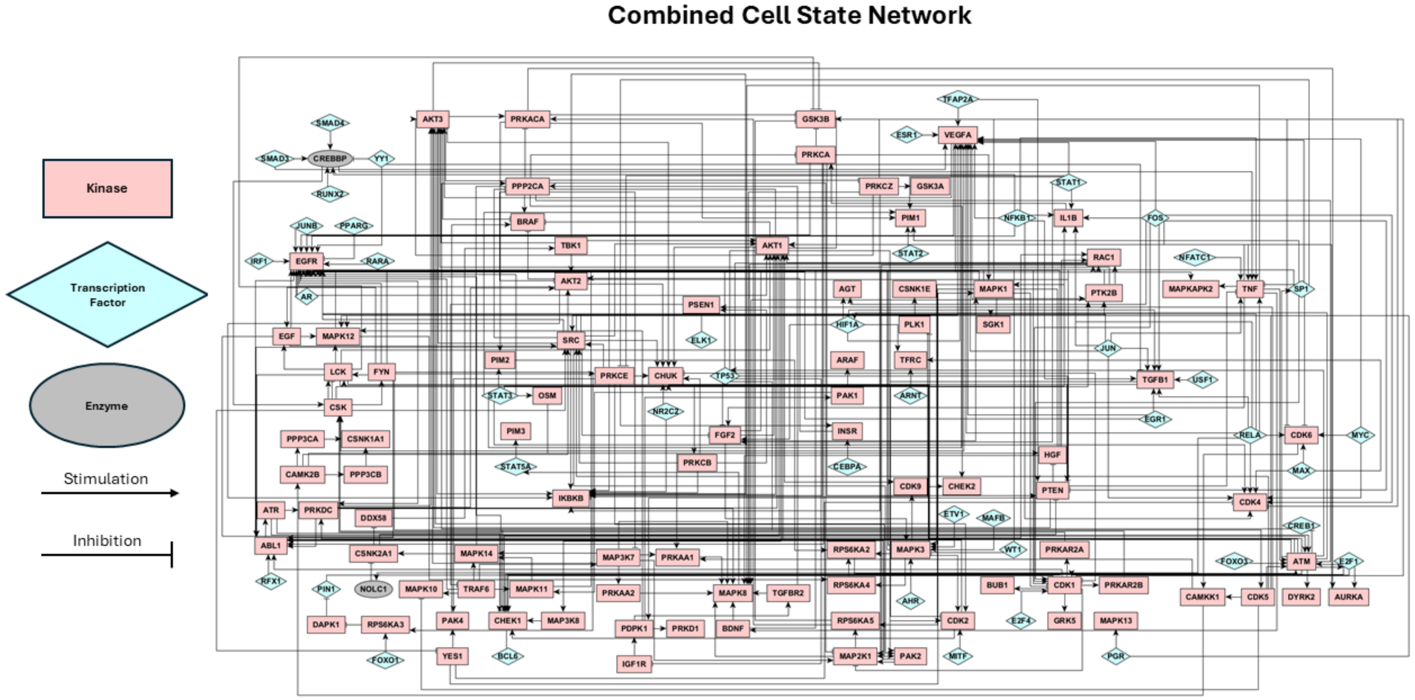
Combined Cell State Network
Kinase
Transcription Factor
Enzyme
Stimulation
Inhibition

**Supplementary Figure 4 (S4). COSMOS PPINs.** The PPINs of the four individual cell states (a-d) and the final combined network (e). Pink rectangles indicate kinases and blue diamonds indicate transcription factors. Activating/Stimulating interactions are depicted by arrows and inhibitory interactions are depicted by T-bars.

#### a) Multinomial Regression

Primary Tumors

| Phenotype | Sensitivity | Specificity | Balanced Accuracy |
|---|---|---|---|
| AC | 1.0<br>1.0<br>1.0 | 0.72<br>0.51<br>0.42 | 86.0%<br>75.5%<br>74.5% |
| MES | 0.97<br>0.73<br>0.71 | 0.99<br>0.99<br>0.95 | 98.3%<br>86.4%<br>83.3% |
| NPC | 0.41<br>0.22<br>0.14 | 0.99<br>0.98<br>0.98 | 70.2%<br>60.8%<br>56.8% |
| OPC | 0.0<br>0.0<br>0.0 | 1.0<br>1.0<br>1.0 | 50.0%<br>50.0%<br>50.0% |

Recurrent Tumors

| Phenotype | Sensitivity | Specificity | Balanced Accuracy |
|---|---|---|---|
| AC | 1.0<br>0.0<br>1.0 | 0.7<br>0.46<br>0.42 | 84.9%<br>23%<br>70.1% |
| MES | 0.98<br>0.64<br>0.6 | 0.99<br>0.9<br>0.96 | 98.5%<br>77.2%<br>78.3% |
| NPC | 0.38<br>0.14<br>0.22 | 0.99<br>0.99<br>0.98 | 68.9%<br>55.1%<br>59.1% |
| OPC | 0.0<br>0.0<br>0.0 | 1.0<br>1.0<br>1.0 | 50.0%<br>50.0%<br>50.0% |

Original Data
Data +10% Noise
Data +20% Noise

#### b) K-Nearest Neighbor (KNN)

Primary Tumors

| Phenotype | Sensitivity | Specificity | Balanced Accuracy |
|---|---|---|---|
| AC | 1.0<br>1.0<br>1.0 | 0.88<br>0.64<br>0.58 | 94.0%<br>82.0%<br>79.0% |
| MES | 0.96<br>0.80<br>0.69 | 0.99<br>0.98<br>0.97 | 98.3%<br>88.7%<br>84.7% |
| NPC | 0.83<br>0.43<br>0.46 | 0.98<br>0.98<br>0.97 | 90.8%<br>70.6%<br>72.0% |
| OPC | 0.0<br>0.0<br>0.0 | 1.0<br>1.0<br>1.0 | 50.0%<br>50.0%<br>50.0% |

Recurrent Tumors

| Phenotype | Sensitivity | Specificity | Balanced Accuracy |
|---|---|---|---|
| AC | 1.0<br>0.0<br>1.0 | 0.82<br>0.62<br>0.57 | 91.1%<br>31.0%<br>78.3% |
| MES | 1.0<br>0.78<br>0.77 | 0.99<br>0.86<br>0.98 | 99.5%<br>82.0%<br>87.4% |
| NPC | 0.76<br>0.32<br>0.35 | 0.99<br>0.99<br>0.99 | 87.8%<br>64.6%<br>66.6% |
| OPC | 0.0<br>0.0<br>0.0 | 1.0<br>1.0<br>1.0 | 50.0%<br>50.0%<br>50.0% |

Original Data
Data +10% Noise
Data +20% Noise

c) **Random Forest (RF)**

Primary Tumors

| Phenotype | Sensitivity | Specificity | Balanced Accuracy |
|---|---|---|---|
| AC | 0.0<br>0.0<br>0.33 | 0.99<br>0.97<br>0.96 | 49.5%<br>48.5%<br>64.7% |
| MES | 0.98<br>0.98<br>0.93 | 0.93<br>0.91<br>0.89 | 95.5%<br>94.4%<br>90.9% |
| NPC | 0.98<br>0.89<br>0.76 | 0.95<br>0.97<br>0.92 | 96.7%<br>93.1%<br>84.0% |
| OPC | 0.0<br>0.25<br>0.25 | 1.0<br>0.99<br>0.96 | 50.0%<br>62.0%<br>60.4% |

Recurrent Tumors

| Phenotype | Sensitivity | Specificity | Balanced Accuracy |
|---|---|---|---|
| AC | 0.0<br>0.0<br>0.0 | 0.95<br>0.91<br>0.92 | 47.3%<br>45.5%<br>45.5% |
| MES | 0.98<br>0.98<br>0.76 | 0.94<br>0.96<br>0.92 | 96.9%<br>98.0%<br>83.6% |
| NPC | 0.98<br>0.92<br>0.78 | 0.99<br>0.99<br>0.97 | 98.5%<br>95.2%<br>87.9% |
| OPC | 0.27<br>0.18<br>0.27 | 1.0<br>1.0<br>0.96 | 63.6%<br>59.1%<br>61.7% |

Original Data
Data +10% Noise
Data +20% Noise

d) **XGBoost**

Primary Tumors

| Phenotype | Sensitivity | Specificity | Balanced Accuracy |
|---|---|---|---|
| AC | 0.0<br>0.67<br>0.67 | 0.91<br>0.86<br>0.81 | 45.6%<br>76.3%<br>73.8% |
| MES | 0.98<br>0.93<br>0.81 | 1.0<br>0.93<br>0.91 | 99.1%<br>93.2%<br>86.1% |
| NPC | 0.89<br>0.76<br>0.70 | 0.95<br>0.98<br>0.94 | 92.3%<br>87.0%<br>82.1% |
| OPC | 0.0<br>0.0<br>0.0 | 1.0<br>1.0<br>1.0 | 50.0%<br>50.0%<br>50.0% |

Recurrent Tumors

| Phenotype | Sensitivity | Specificity | Balanced Accuracy |
|---|---|---|---|
| AC | 0.0<br>0.0<br>0.0 | 0.83<br>0.76<br>0.74 | 41.6%<br>38.0%<br>37.1% |
| MES | 0.98<br>0.80<br>0.75 | 0.99<br>0.78<br>0.92 | 98.2%<br>78.7%<br>83.6% |
| NPC | 0.81<br>0.51<br>0.67 | 0.99<br>0.94<br>0.91 | 89.8%<br>72.4%<br>79.2% |
| OPC | 0.0<br>0.0<br>0.0 | 1.0<br>1.0<br>1.0 | 50.0%<br>50.0%<br>50.0% |

Original Data
Data +10% Noise
Data +20% Noise

**Supplementary Figure 5 (S5). Machine Learning Results.** The sensitivity, specificity, and balanced accuracy for each machine learning model. Results are divided into two tables, one for primary tumors and one for recurrent tumors. Results from the original GLASS data is shown in red, while results from the data with 10% and 20% noise are shown in light and dark blue, respectively.

### ADDITIONAL SUPPLEMENTARY FILES

**Additional File 1: Differentially_Expressed_Genes.xlsx**. This file contains the lists of differentially expressed genes in each of the four dominant cell states of glioblastoma. Each cell state is a separate sheet in this order: MES, NPC, OPC, AC.

**Additional File 2: Differentially_Expressed_Phosphoproteins.xlsx.** This file contains the lists of differentially expressed phosphoproteins in each of the four dominant cell states of glioblastoma. Each cell state is a separate sheet in this order: MES, NPC, OPC, AC.

**Additional File 3: COSMOS_PPI.xlsx** This file contains information regarding the PPINs for each of the four dominant cell states, as well as the final integrated network (5 total). Each network has two sheets associated with it. Sheets with the suffix "ATT" describe the attributes of the nodes of the PPIN, including the protein name and activity. Sheets with the suffix "SIF" describe the interaction between nodes, which is whether the interaction is activating or inhibiting.
